# Ordinary differential equations to construct invertible generative models of cell type and tissue-specific regulatory networks

**DOI:** 10.1101/2023.05.18.540731

**Authors:** Eliatan Niktab, Paul H. Atkinson, Mark Walterfang, Ingrid Winship, Stephen L. Sturley, Andrew B. Munkacsi

**Affiliations:** Applied Technologies, Callaghan Innovation, WLG, New Zealand; School of Biological Sciences, Victoria University of Wellington, WLG, New Zealand; Maurice Wilkins Centre, Metabolic Health, AKL, New Zealand; Neuropsychiatry, Royal Melbourne Hospital, Melbourne, Vic, Australia; Department of Psychiatry, University of Melbourne, Melbourne, Vic, Australia; Florey Department of Neuroscience and Mental Health, Melbourne, Vic, Australia; Department of Genomic Medicine, Royal Melbourne Hospital, Melbourne, Vic, Australia; Department of Medicine, University of Melbourne, Melbourne, Vic, Australia; Department of Biology, Barnard College, Columbia University, NY, USA

## Abstract

Single-cell RNA-Seq (scRNA-seq) transcriptomics can elucidate gene regulatory networks (GRNs) of complex phenotypes, but raw sequencing observations only provide ”snap-shots” of data and are inherently noisy. scRNA-seq trajectory inference has been utilized to solve for the missing observations, but disentangling complex dynamics of gene-gene interactions at different time points from aggregated data is a non-trivial task and computationally expensive. Here we describe our Non-Stiff Dynamic Invertible Model of CO-Regulatory Networks (NS-DIMCORN) to define the genetic nexus underpinning specific cellular functions using invertible warping of flexible multivariate Gaussian distributions by neural Ordinary differential equations. Our results yield a generative model with unbiased density estimation from RNA-seq read-count data only. This resulted in scalable time-flexible sampling of each gene’s expression level thence allowing ab initio assembly of gene regulatory networks in specific cells. We demonstrate our proposed methodology is superior to the state-of-the-art algorithms in accurately recovering genome-wide functional interactions, whether from synthetic or empirical data. We optimized our algorithm for GPU-based implementation thereby further enhancing the utility of our proposed methodology in comparison to the ten benchmarked methods.

## 2 Introduction

Historically, GRNs are represented where genes are nodes, edges are pairs of genetic interactions, and networks can be assembled from overlapping pairs. GRNs may be inferred from gene deletion (perturbation) screens and experiments that compared non-perturbed (healthy) versus perturbed (diseased tissues)^1^. More recently emergence of high-depth multi-sample (bulk RNA-seq) and single-cell RNA-sequencing (scRNA-seq) data allowed more accurate GRN inference by modeling dynamic RNA expressions for different tissue types or cells at different developmental stages (time points)^2^. Distilling informative *ab initio* genome-wide GRNs from whole-genome RNA expression is especially useful given that Next Generation Sequencing (NGS) can conveniently achieve statistical power using pseudo-bulk techniques and high-depth single-cell RNA sequencing^3^. NGS has already produced substantial tissue and cell-specific RNA-seq data that would allow comparing genetic interactions in different experimental settings (e.g., healthy/diseases), different cellular processes, and different developmental stages, all at tissue/single-cell level resolution^4^.

In RNA-seq experiments, the expression level of transcripts is calculated from the number of sequenced reads that map to the codon responsible for those transcripts with no time stamping^5^. Thus, Differentially Expressed (DE) transcripts can also indicate the direction and strength of their correlation with other genes and samples^6^. However, it should be noted that RNA-seq data from NGS contains high technical noise, which can be exacerbated by sequencing-specific data features such as sample heterogeneity, variation in sequencing depth, and sparsity mapped reads^2^. Additionally, regulatory interactions are deemed far more interconnected than simple loci-loci interactions due to the discovery of various small Ribonucleic Acid (RNA) sequences that play active roles in the machinery that regulates cellular processes^7^ which must be taken into account in determining GRNs. Albeit, as reviewed by Pratapa *et al.,* constructing informative models that recapitulate the complete and accurate set of genetic interactions from RNA-seq data is now a very active area of research^8^.

Initial *ab initio* GRNs based on Boolean logic^9^ have been successfully used to model high-level monotonic interactions of cellular mechanisms^10^. Even so, these models failed to capture cascades of complex events, such as promoter recognition and the dynamic self-regulatory protein translations across the entire genome, over the differentiating lifetime of a cell^11^. Therefore most recently, enhanced Boolean logic^12^, regularized linear regressions^13^, Bayesian networks^2, 14^, partial correlation, semi-partial correlation^15, 16^, Pearson correlation^17^, tree^11^, entropy^18^ based approaches^19^ and Gaussian graphical models^20^ have all been utilized to model intricate interactions between genes and gene products (e.g., proteins), but accurate GRNs inference remains a challenging problem^8^. Briefly, enhanced Boolean logic, regression, and correlation-based approaches fail to capture higher-order and more complex gene-gene relationships (e.g., non-additive interactions)^9^. Though such interactions can be elucidated using mutual information entropy, they require homogeneous data or hyper-parameter re-tuning to avoid overestimating interactions’ significance^21^. Spanning tree-based approaches can be used but are computationally expensive, extremely sensitive to changes in data, and inadequate for predicting continuous values as observed in empirical data^22^. Traversing Bayesian networks may also be used, but calculating the conditional probability of edges is not a trivial problem when a large number of interactions are involved, and these networks are only suitable for steady-state data in their vanilla form^2, 14, 20^. Finally, Gaussian graphical methods depend on the Gaussianity assumption, which also implies linear dependencies between genes; additionally, most implementations of this approach to date are incapable of handling directional interactions as in cell differentiation^16^.

Another promising approach for inferring GRNs from time-stamped data (like RNA-seq) is borrowed predominantly from other fields of natural sciences (e.g., physics) that have a long history of studying dynamic systems that lend themselves to the dynamic processes of biological cells using systems of Ordinary Differential Equations (ODEs)^23^. Such approaches have been demonstrated to generate more realistic behavior and better describe genuine regulatory relationships of genome-wide interactions in cells^4, 20^. In ODE-based methods, derivatives of differential equations describing the system can be estimated by difference approximation^24, 25^ or regularised differentiation^26^, and then be solved by linear methods^27^ by fitting a mechanistic of nonlinear functions^23, 24^ or nonparametric techniques^28^. In both of these approaches, the main constraint of solving ODEs for a biological system with tens of thousands of genes is that such high-dimensional parameter spaces require larger than number of dimensions, sample sizes that can rapidly become computationally intractable as the addition of new samples adds up^2^. Here we demonstrate that this problem can be dealt with satisfactorily by

1. utilizing a highly flexible and scalable method that can model functional relationships of dynamic data
2. bypassing the error-prone derivative estimation, and
3. using a nonrestrictive and scalable model architecture that allows cheap computation.

We describe the Non-Stiff Dynamic Invertible Model of CO-Regulatory Networks (NS-DIMCORN) that allows unrestricted neural network architectures (i.e., arbitrary depth increases) and training the model without partitioning or ordering the data dimensions. NS-DIMCORN yields a continuous-time invertible generative model with unbiased density estimation by one-pass sampling, allowing scalability and end-to-end training of larger ODEs-based models (Figure 1). Furthermore, NS-DIMCORN only requires scRNA-seq read count data as an input, and time points are automatically inferred by estimating probability distributions for the continuous gene expression trajectories instead of probability distributions for the derivatives. This allows easy sampling of the continuous trajectories using Hamiltonian Monte Carlo and calculates nonlinear gene dependency based on conditional Mutual Information (MI)^29^. To this end, we demonstrated that NS-DIMCORN, on average, outperforms nine other state-of-art algorithms^15, 17–19, 25, 30–33^ in inferring GRN from synthetic, bulk, and single-cell RNA-seq data.

**Figure 1.**
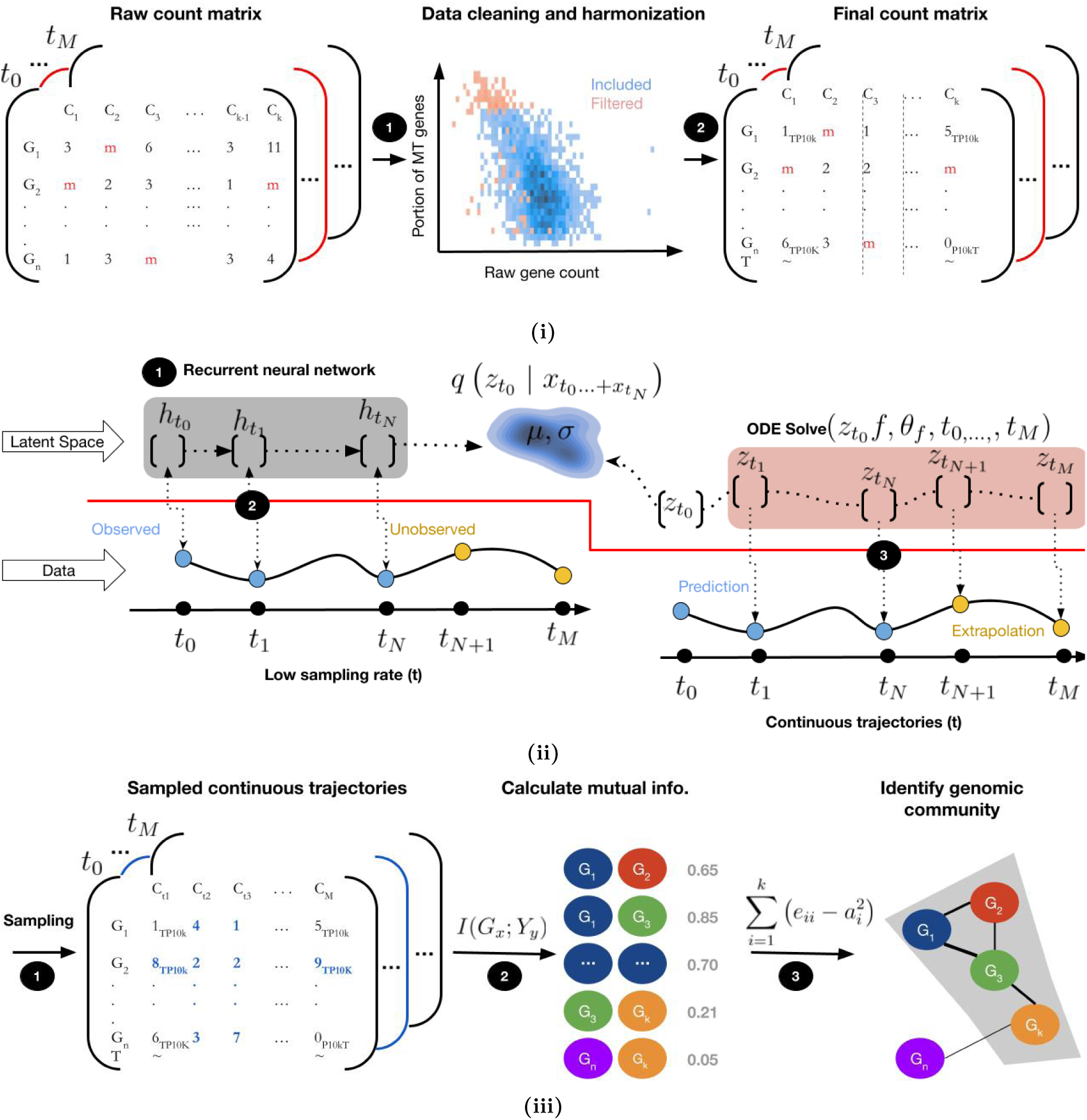
Overview of NS-DIMCORN. i: Data cleaning and harmonization where mitochondrial genes (MT), cells with low gene count, batch effects, and genes with low depth were removed. ii: training a neural network for bijective mapping from a latent trajectory of learned continuous dynamics to data. iii: sampling and constructing sample-specific GRN’s for community detection, from TP10K as defined at Section 4.2.

## 3 Results

### 3.1 Overview of the algorithm

Given a set of irregularly sampled (missing reads indicated in red in Figure 1i) scRNA-seq data for a specific tissue or cells, the goal of NS-DIMCORN is to model gene expression across cellular process trajectories (i.e., cell lineage differentiation trajectories). To this end, the read counts for all the scRNA-seq samples are normalized (Figure 1i-1); samples with spurious or low-quality reads are removed; counts are standardized, and the data set is harmonized to remove any confounding variables such as batch effects (Figure 1i-2). NS-DIMCORN represents different cell states by a continuous latent trajectory (Figure 1ii-1) and defines a bijective map from the latent learned latent space to data by integrating latent variables (Figure 1ii-2). Latent trajectories are computed by solving an initial value problem by an ODE solver that is parameterized by a neural network and a given initial state, *z_t_*_0_. The output of the last layer of the neural network is the solution to the initial value problem, the hidden units are parameterized as a continuous function of time, and the parameters of nearby “layers” are automatically tied together. This learned model then, in turn, allows continuous sampling of reading counts even for missing states/genes (blue states and reads) arbitrarily far forwards or backward in time (Figure 1iii-1). Continuous sampling from cell states allows accurate estimation of conditional mutual information between each set of gene pairs as *ab initio* (translated to weights of edges between two genes) and signs of co-variances as an indicator of interaction directions in the inferred networks of tissue-specific gene-gene interactions (Figure 1iii-2).

**Figure 2.**
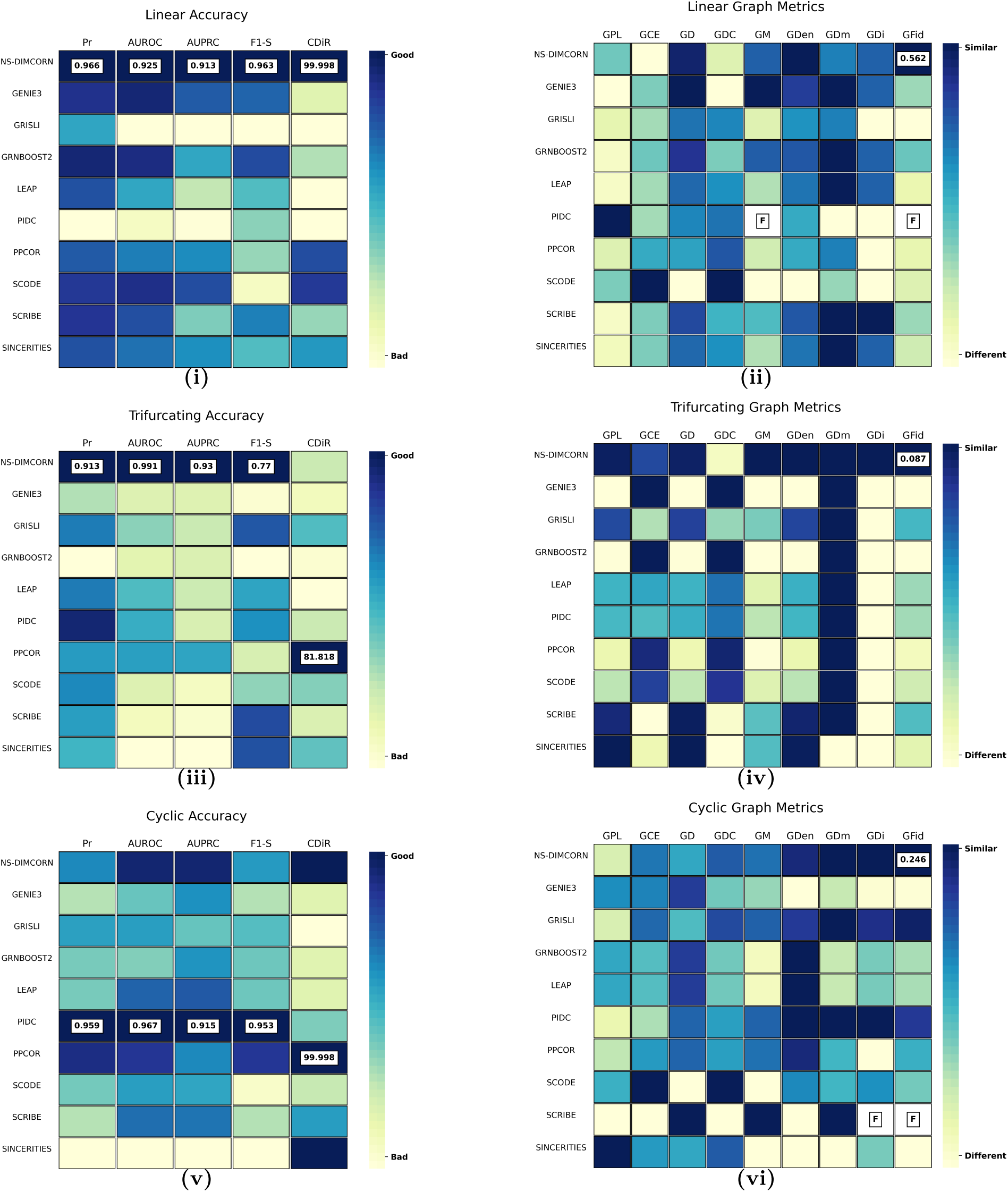
Comparison of different algorithms when inferring GRNs from synthetic data. In i, iii, and v, the six measurements (columns) represent a measure of accuracy (as described in the methods section) for each algorithm (rows). In ii, iv, and vi Absolute Minkowski distance of eight fundamental graph metrics between the ground truth network and inferred network and the Jensen-Shannon divergence of Laplacian spectra of the two graphs (GDi) are plotted. Column values for are panels are normalized, where blue represents better accuracy/faithfulness than yellow. Blocks with the highest values and more than three significant figures of difference are annotated, and F indicates an undefined number. Overall, the heatmaps presented show NS-DIMCORN stays faithful to the ground truth network topology and demonstrates the best accuracy for simulated long linear and trifurcating development processes. PIDC slightly outperforms NS-DIMCORN in the sense of accuracy by less than 1 percent but NS-DIMCORN constructed the most accurate GRN topologically.

### 3.2 Benchmarking against simulated data

#### 3.2.1 Simulated data for Long Linear (Li), Trifurcating (Tr), and Cyclic (Cy) cellular processes trajectories

In a closed dynamic system, functional changes result from the interchange of information and interactions between constructional elements of the system (i.e., structural elements can switch each other on/off or self-regulate to steer the fate of that system)^34^. In the context of GRNs, nuanced and continuously evolving changes of these dynamic systems can be surveyed employing network topological features, which can elucidate the type and strength of pairs and groups of interactions between genes in a network^35^. Many metrics have been suggested to study a network’s topology and estimate the structural distance (similarity) between two networks^36^.

Here we focused only on the most robust and commonly used measures of network topology^37^, namely Graph Path Length (GPL), Graph Degree (GD), Graph Modularity (GM), Graph Diameter (GDm), and Graph Clustering Coefficient (GCE) as defined in the Methods (Section 4.9.1). For ease of reporting, we equipped the vector space of these metrics with a norm–as opposed to using the topology metrics directly. We thus reported the Minkowski distance of each one of these metrics for the inferred network and their respective Ground Truth Networks (GTNs) in both synthetic examples and Known to be Truth Sub-Networks of experimental data (KTSNs). Our analyses only included interactions experimentally validated interactions for empirical data (Section 4.8). In addition to topology metrics, it has been demonstrated that the Laplacian eigenvalues of graphs also capture the local and global properties of networks^38^; therefore, we further included Graph Distance (GDi) metrics which capture the structural distance of the compared networks in terms of the Jensen-Shannon distance of graph Laplacian matrices (Section 4.9.6). Finally, combining all the proposed metrics, we calculated Graph Fidelity Distance (GFid) that normalizes and averages measurements in more than one graph (Section 4.9.7)^37^. Using these considerations, we compared ten tools for inferring *ab initio* GRNs, including NS-DIMCORN. We sought to determine if GRNs inferred by an algorithm from a database would exhibit dynamics identical to the known underpinning network verified as described in (Section 4.8.2). To this end, we executed each algorithm on pre-processed expression data as described in Methods using the recommended hyper-parameters from the original publications^15, 17–19, 25, 30–33^. This step resulted in a ranked edge list of the inferred network connections (edges). After removing all the self-loops, the inferred networks were analyzed for faithfulness to the true network and accuracy as described in the Methods (Sections 4.8 to 4.9), we then reported two sets of statistics indicating the performance of each of the cited algorithm.

To avoid the pitfall of technical and instrumental noise in scRNA-seq data^8, 39, 40^, and for ease of interpretation^41^, we first focused on synthetic data with a known GRNs that could serve as the ground truth (Figures 2 to 3). This initial focus also allowed us to avoid any limitations of pseudo-time dependent inference that could potentially affect our benchmarking results, especially for GRISLI^30^, LEAP^17^, PPCOR^15^, SCODE^32^, SCRIBE^33^, and SINCERITIES^19^ which required pseudo-times provided separately as input. Three different temporal trajectories here, namely Linear, Cyclic, and Trifurcating, are constructed as described in BoolODE package^42^ represent the different possible dynamic cellular processes^8^, whether the trajectories relate to metabolism, cellular reprogramming, reproduction, differentiation, or apoptosis through cell developmental stages. As for the linear trajectory here, we included a long cascade of intermediate genes to attain enough complexity but ensured the linear trajectory still resulted in one distinct final steady state for each initial state.

**Figure 3.**
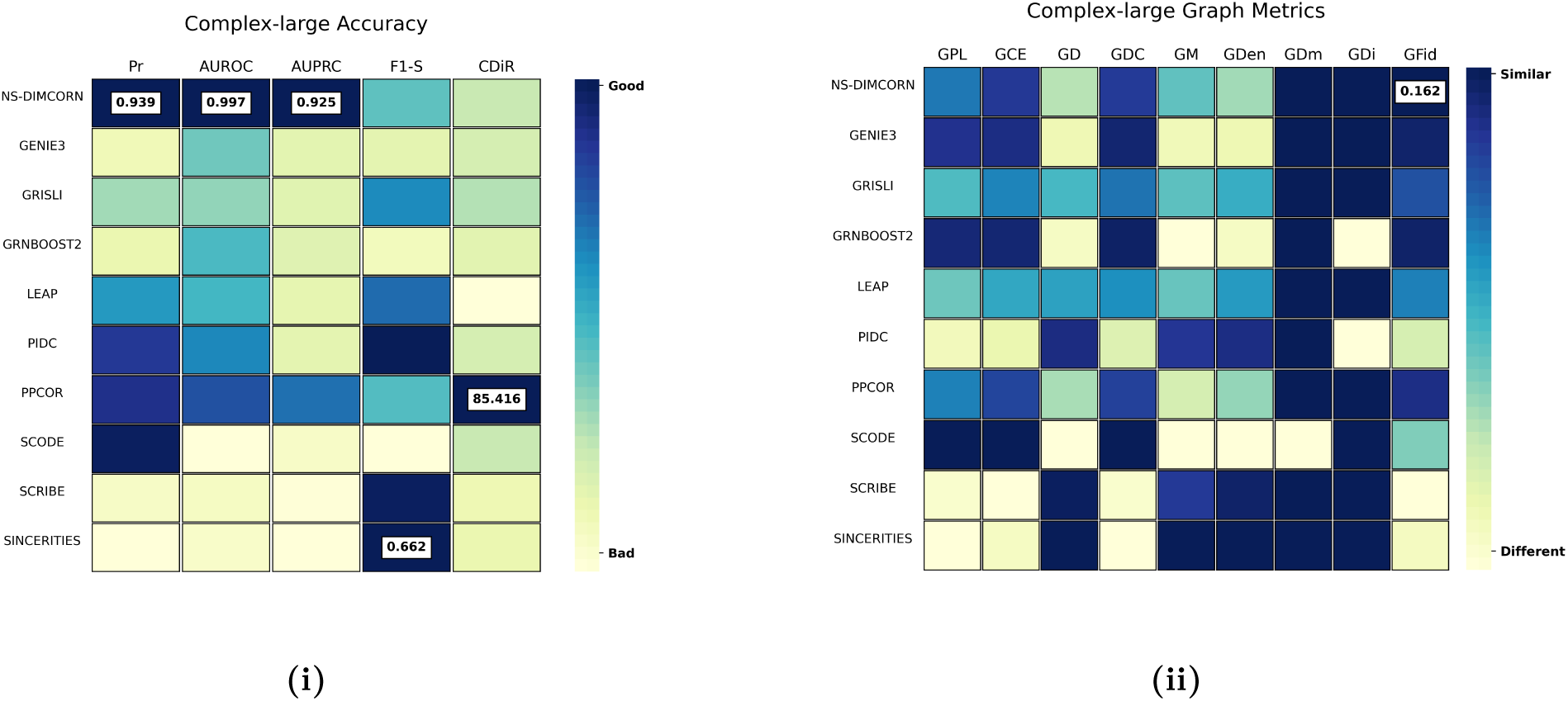
Comparison of different algorithms inferring GRNs from cells with complex dynamics (linear, trifurcating, and cyclic) with large sample sizes and a high number of active genes dynamic.

We observed that GRNBOOST2^31^ and SCODE identified the highest number of genuine regulatory interactions for long linear trajectories. At the same time, PIDC failed to identify any interaction owing to its approach in calculating mutual information between genes. However, GRNBOOST2 and SCODE, notably SCODE, showed low discrimination power, resulting in many false positives, spurious interactions, and consequently lower AUROC and AUPRC –indicators of discrimination power (Figure 2i). For the same linear trajectory, NS-DIMCORN only misclassified one of the authentic regulatory interactions (Section 4.8.2) but with far fewer false positives than other methodologies and inferred the most accurate regulatory network, with the highest AUROC and AUPRC. The non-perfect Direction scores (CDiR) for NS-DIMCORN also resulted from some misclassified genuine interactions. Otherwise, the correct direction was inferred for every identified genuine interaction (Figure 2i). Expectedly, topology analysis of the inferred graph confirmed the above model accuracy metric, and NS-DIMCORN showed the highest fidelity toward the ground truth. We hypothesized that the unanticipated high difference in GCE (see Methods for metric definitions) for NS-DIMCORN can be attributed to the fact that the clustering step of NS-DIMCORN creates artificial groupings within the harmonized structure of the linear trajectory and hence reduces the statistical power of the NS-DIMCORN inference. Indeed, omitting the clustering step improved the performance of the NS-DIMCORN for the linear trajectories (Supplementary Table S1), but we believe a larger sample size would be a less error-prone approach for resolving this issue, primarily when cell trajectories are not known or suspected to be complex (Figure 2ii).

We also analysed trifurcating trajectories of cellular processes where mutual regulation motifs involving more than one gene result in a few distinct steady states from common initial states. As illustrated in Figure 2iii, benchmarking NS-DIMCORN against the nine other algorithms for trifurcating trajectories demonstrated the highest precision and superior discrimination power of NS-DIMCORN, based largely on the highest AUROC (Section 4.7.3) and AUPRC (Section 4.7.2). Nonetheless, NS-DIMCORN was mostly unsuccessful in identifying the direction of trajectory interaction. This was because of the arbitrary Cartesian coordinate of each final steady state. Indeed NS-DIMCORN intrinsically would not have any notion of the origin and would also not capture this information in the latent space. PPCOR, on the other hand, was able to identify directions better, although for the fewer correctly identified genuine interactions, using partial direction correlation between two pairs and relying on the direction of pseudotimes (Figure 2iv).

NS-DIMCORN had difficulty with oscillatory circuits that yield linear trajectories where the final state coincides with the initial state. This behavior was mapped out here in the synthetic cyclic data with zero steady-state. NS-DIMCORN was only the second-best inference algorithm based on AUROC as well as AUPRC and demonstrated lower precision than PIDC and PPCOR. Circularity introduced by the absence of initial/final temporal distinction among cells most likely underlies this lack of performance. Regardless, NS-DIMCORN still successfully captures the actual topology of the original graph better than all the other methods studied here (Figure 2v).

#### 3.2.2 Simulated data for Compound-Complex (CC) cellular processes trajectories

We did not expect that the relatively simple rules used by BoolODE to generate synthetic data, even in their complex mode, would sufficiently mimic the properties of the real biological data. So we sought to determine if a GRN inferred by an algorithm from an empirical dataset would exhibit dynamics and steady states identical to the original underpinning network. Empirical RNA-seq data is often categorized into three distinct groups (steady-state, bulk, and single-cell sequencing data)^43^. The steady-state data refer to the expression level of genes after introducing gene knockouts or essential gene perturbation under the assumption that only meaningful pairwise interactions exist^44^. Even after excluding non-characterized transcriptomes, this approach requires more than 200 million observations to assay all pairs of genetic interactions for the remaining *≈* 20, 000 human genes^45^. At the time of writing that paper, the largest dataset of this type only included around 0.1% gene pairs^46^ of possible interactions.

The other two types of data, namely Bulk RNA-seq data and scRNA-seq data, provided sample data sets comprising snapshots of different cellular process states within different cells or tissues^47^. Consequently, these data can represent the dynamic interactions of genes and cellular process trajectories beyond genetically modified or chemically perturbed, which is the focus of this study^43^. While scRNA-seq data does not suffer the loss of information inherent to averaging processes in bulk RNA sequencing methodology^47^, it is more noise-prone^48^.

To establish that NS-DIMCORN is scalable but at the same time is also sensitive enough to detect nuanced dynamics specific to different tissues in the human body among the noise, we included a larger dataset of brain cells with more specialized cell types and a smaller dataset of heart cells. Empirical ground truths mainly rely on prior expert knowledge and, due to their binary and agglomerative nature, do not encapsulate signal type, direction, or time dependency of the functional genomics nexus they describe^49, 50^. Moreover, pleiotropic effects of genes and noisy high throughput data used for the curation of these networks, combined with the subjective allocation of genes to a network, cause truth networks to differ in number and type of genes^51^ in different studies and may include many spurious interactions. To alleviate the effects of false positive or false negative interactions from empirical ground truth data sets in our experiments, we constructed a ground truth network comprising only experimentally validated Transcription Factors (TFs) and genetic interactions that are involved in essential and well-characterized metabolic pathways (Section 4.1).

#### 3.2.3 NS-DIMCORN inferred network from sc-RNA-seq data is cell specific

In the liver, SCODE identified the highest GII (Section 4.8.2) compared to its respective KTSN (Section 4.1.2) and many other interactions. The high number of interactions identified might indicate many false positives; therefore, a worse T5KI (Section 4.8.4) score was given to SCODE (Figure 4i). PPCOR more or less showed the same behavior as SCODE, which is in contrast to NS-DIMCORN, which attained a better GII score and showed good sensitivity indicated by low AII (Section 4.8.1) and high T5KI (Section 4.8.4). Comparing the topological fidelity distance (Section 4.9.7) of the KTSN and the inferred GRN again demonstrated the superior performance of the NS-DIMCORN. GM (Section 4.9.3) and GD (Section 4.9.6) scores are also consistent with our rationale that SCODE connected most of the genes in the network and lost the modular texture of real GRNs (Figure 4ii).

**Figure 4.**
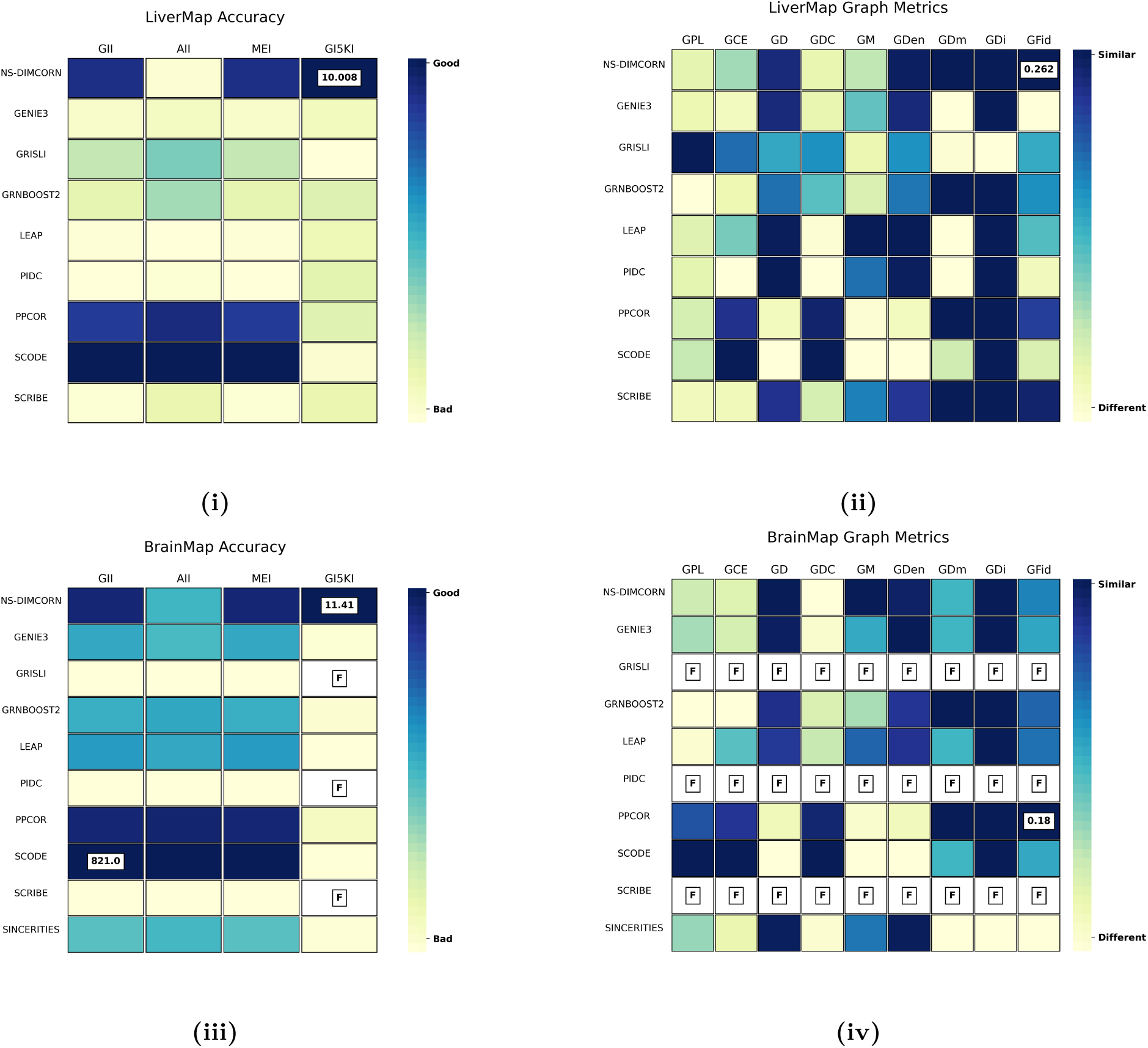
NS-DIMCORN is scalable and identifies more known genuine metabolic interactions than any other algorithm in brain and liver single-cell sequencing data. a;c, The four measurements (columns) represent a measure of accuracy for each algorithm (rows) where only true positives are partially known. b;d, Absolute Minkowski distance of eight fundamental graph metrics between sub-networks of partially known genuine metabolic interactions and inferred networks in addition to Jensen-Shannon divergence of Laplacian spectra of the two graphs (GDi) are measured. Column values are normalized, darker blue represents better accuracy, and lighter yellow represents lower accuracy. Blocks with the highest values and more than three significant figures of difference are annotated, and F indicates an undefined number.

In the brain, NS-DIMCORN again achieved the highest GI5KI but showed a more significant topological distance to the KTSN than PPCOR. Graph path length (Section 4.9.1) and gene centralities appeared to contribute most to this observation (Figure 4iii, Figure 4iv). The extra interactions identified by PPCOR appear to be non-overlapping with the KTSN, but the same genes, in general, having more interactions as indicated by graph clustering coefficient. The greater reach regardless of their authenticity would explain this observation. It is noted that GRISLI, PIDC, and SCRIBE failed to generate an output, given the size of the brain dataset and the number of genes involved in the network. GRISLI ran out of memory on a high-performance 72-core computer with more than 2TB of RAM, SCRIBE could not generate output in more than seven days on the same computer, and PIDC produced only undefined values for the output (Figure 4iii, Figure 4iv).

#### 3.2.4 NS-DIMCORN is scalable and allows specificity for bulk RNA sequences and aggregated data

For the bulk RNA-seq data, despite the fact that the brain has more specialized cells, only a few people with different sub-brain tissue information were obtainable. Liver samples were of a larger dataset, although the number of individuals sequenced for the liver data was still relatively small compared to the number of cells in the scRNA-seq dataset. Our rationale for choosing bulk RNA-seq data was to investigate if NS-DIMCORN can still successfully infer the best network when the data is highly aggregated–bulk, many genes are involved–the brain or only partial data exist–few sub-tissues. Our observations were largely in accordance with the result from scRNA-seq data. The only difference is that we recovered a more accurate topology for the bulk brain, which is consistent with the literature stating bulk RNA might still be a better choice for identifying interactions with smaller effect sizes^47^. We also observed SCRIBE was the only algorithm that failed on the brain data, which suggests, as opposed to GRISLI and PIDC, the algorithm performance is dependent on the network size (Figure 5iv).

**Figure 5.**
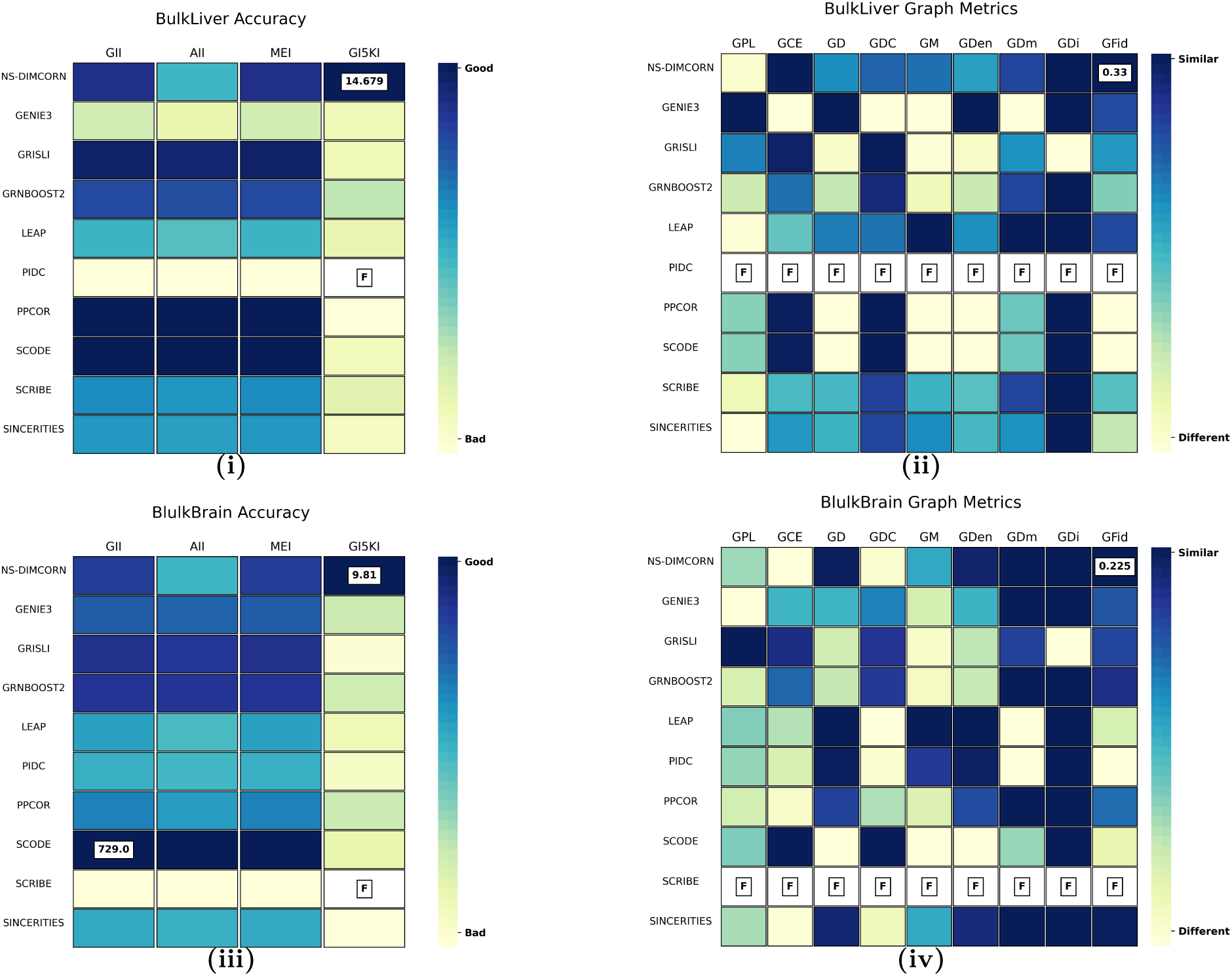
NS-DIMCORN is less sensitive to diffident cell types in the same tissue in comparison to different but similar tissues.

## 4 Discussion

We excluded four algorithms that we included initially in this study, namely SCRIBE^33^, SINGE^52^ GRN-VBEM^14^ and SCNS^12^ given that they failed to produce an output for most or all of the datasets studied here. Speed, memory, or inflexible implementation were the main drawbacks of these methods on our high-performance computer with 2TB of RAM, 72 core CPUs, and 4 Nvidia A100-80GB GPUs. Although these specs are vastly better than an ordinary desktop computer, not every method could achieve concurrency or efficiently utilize all the provided computation resources, so the same result would likely have been obtained on low-spec computers. We did not attempt to optimize the run time of any of these methods and terminated any process after a week if no output was produced. Of these methods, NS-DIMCORN was the only method cable of utilizing GPUs for array operations and model training, being entirely based on TensorFlow^53^ and cuPy^54^ (a GPU based implementation of NumPy). We observed that including 2500 genes for around 8000 cells during training, the NS-DIMCORN model requires roughly thirty hours to model the data. However, our method’s run-time and memory usage heavily depend on the number of genes and samples used as input.

NS-DIMCORN primarily relies on DESC for clustering the related cells before training the generative model. We note, however, this might introduce some limitations when studying linear cellular trajectories. DESC originally initialized clustering regions using the Leiden clustering algorithm^55^ and then optimized the clusters by stochastic gradient descent^56^. Here we swapped Leiden clustering for DensMAP, believing UMAP better preserves the fine details of the data manifold^55^; therefore, more lenient clustering configurations for UMAP would improve NS-DIMCORN performance for linear data^57^.

Regulatory interaction between genes in different cells can be statistically defined by information the-ory^21^. Although most gene pairs satisfy linear or monotonic relationships, Mutual information (MI) is often used as a generalized correlation measure^58, 59^. It has been suggested that bi-weight mid-correlation transformed via the topological overlap transformation is a more robust correlation measure and attains better accuracy in identifying interacting gene sets in terms of Gene Ontology enrichment^58^. We believe this assumption naively overlooked the implicit reliance of the used MI methodology on the local uniformity of the underlying joint distribution, which is not the case for strongly dependent variables such as gene expression level^18^. NS-DIMCORN uses covariance to identify genes interaction direction and Non-parametric entropy estimation to compute the degree of this correlation. The basic idea behind non-parametric entropy estimation is locally estimating the log probability density at each data point, then averaging these estimates, thence allowing accurate estimation of MI between two strongly dependent variables^60^.

In summary, we presented the Non-Stiff Dynamic Invertible Model of CO-Regulatory Networks (NS-DIMCORN) and systematically evaluated NS-DIMCORN with synthetic data representing different cellular trajectories, bulk, and scRNA-seq data from different tissues and sample sizes. We demonstrated NS-DIMCORN is scalable due to its unrestricted neural network architectures and showed its superior performance compared to other state-of-the-art algorithms for *ab initio* GRN inference based on cellular trajectories. We showed that not only does NS-DIMCORN estimate the chronological ordering of the cellular trajectories unsupervised from the data, but it also offers high sensitivity and specificity due to its invertible generative model that allows unbiased density estimation using continuous sampling.

### 4.1 Datasets

#### 4.1.1 Empirical data

For single-cell sequencing data, the Allen brain map dataset^61^ comprising 76,533 total nuclei from the primary motor cortex of two coronal post-mortem human brain specimens was chosen. Those authors have described details of performed DNA-seq sequencing and preliminary data processing steps such as case inclusion criteria, nucleus dissociation/sorting, and RNA-sequencing methodology (barcode extraction, mapping, alignment, filtering and annotating BAM file with gene tags)^61^. We obtained the raw gene expression count matrix as a CSV file and applied filtering and clustering as described later in (Section 4.2). To study the tissue specificity of our method, we also included the Human Protein Atlas dataset^62^ that comprised 8439 total nuclei derived from parenchymal and non-parenchymal of fresh hepatic tissue of five human livers^63^. We obtained the raw gene expression count matrix that was prepared as described^63, 64^ and applied filtering and clustering specific to this study (Section 4.2). Bulk RNA-seq data in this study was obtained from the GTEx Consortium atlas^65^ portal (dbGaP Accession phs000424.v8.p2). For all the individuals for which data from the liver and brain was available, we downloaded read counts for the RNA-Seq data and processed the data as described in more detail here (Section 4.2).

#### 4.1.2 Known to be true sub-network of empirical data (KTSN)

We only included experimentally validated transcription factors (TFs)^66^ and genetic interactions involved in essential and well-characterized metabolic pathways^67, 68^ in our ground truth as described earlier. We further filtered interactions of the network to “super pathways” only if they have been captured analogously in KEGG^67^, Reactome^68^ and WikiPathway^69^, the three most cited datasets in published -omics studies^51^ to indicate biological evidence.

#### 4.1.3 Synthetic data

Expression data were simulated for 500 cells following linear, cyclic, or trifurcating two-dimensional projections by converting their Boolean GTN interaction matrix into noisy nonlinear ordinary differential equations described by Pratapa et al.^8, 70^ and elsewhere^42^. Random Gaussian noise^71^ was added to ensure the intrinsic stochasticity of the data was conserved in the simulated data^70^.

### 4.2 RNA-seq data prepossessing

We obtained the count matrix (Allen brain, Human Protein Atlas Dataset) and OMNI SNP Array Intensity files (GTEx brain and liver) and then read those files into an AnnData object with hierarchical data format^72^ for the downstream processing. Genes detected in less than a threshold number of cells/tissue samples due to the low sampling rate were not included This threshold was determined based on the average sequencing depth for each dataset so that the majority of high-confidence reads were retrieved (Allen Brain dataset = 100 cells, Human Protein Atlas dataset = 50 cells, GTEx brain = 25 samples, and GTEx liver = 10 samples). To further remove poor-quality cells, we calculated the total RNA read counts and the percentage of those counts relating to mitochondrial genes and then removed cells and samples without enough RNA reads ( 3 *×* Mean Absolute Deviation (MAD)), gene coverage ( 4 *× MAD*), or a high portion of mitochondrial RNA (3 *× MAD*) (Figures 6 and 7).

**Figure 6.**
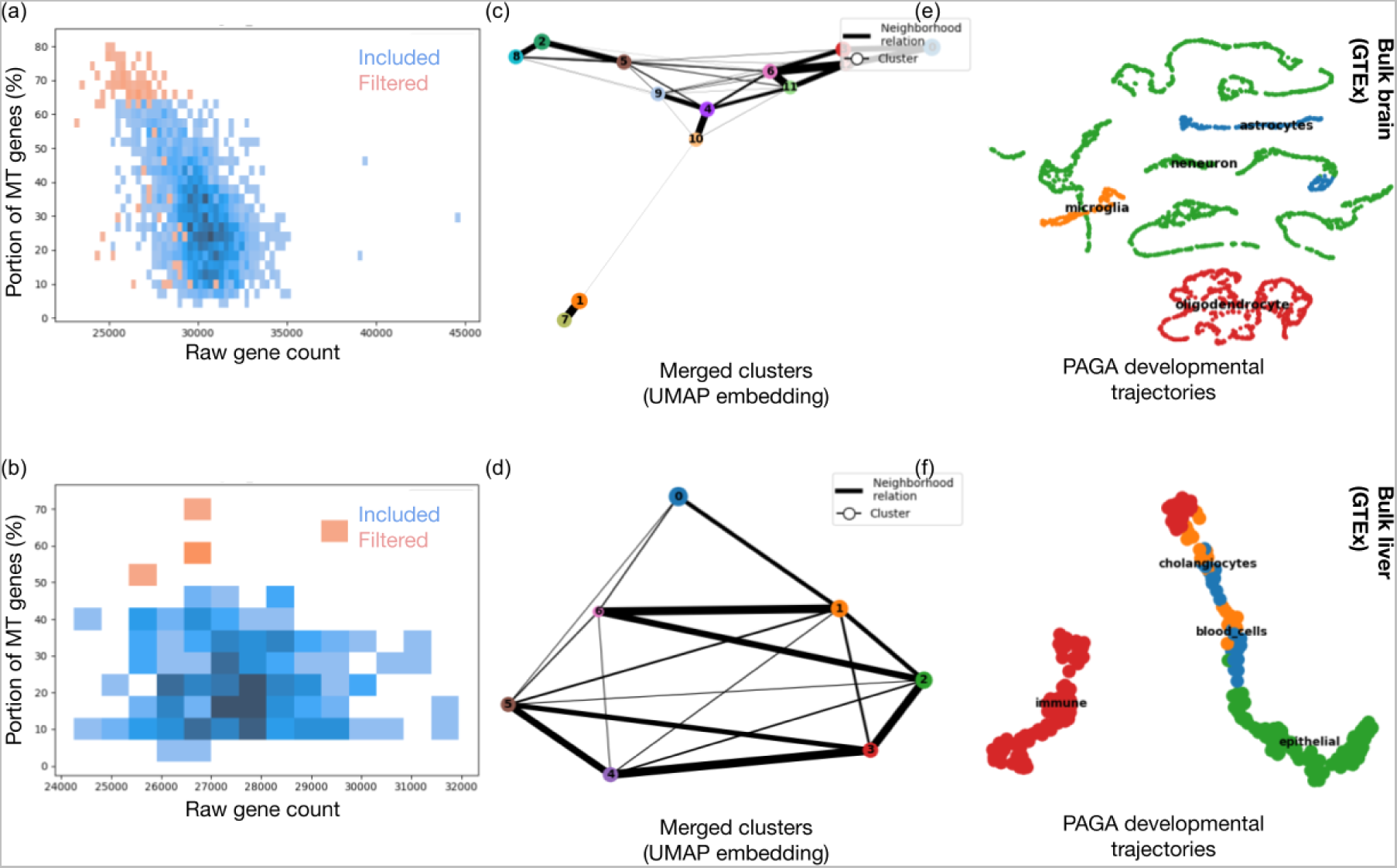
Bulk RNA-seq data prepossessing: cells with less than a threshold RNA-seq reads count or cells with a high portion of mitochondrial genes were removed (a,b). Cells with high-quality reads were then clustered on the expression profile, and PAGA developmental trajectories were calculated (c, d). Obtained clusters were refined into merged sets of the same developmental stage and cell subtype(c, d).

**Figure 7.**
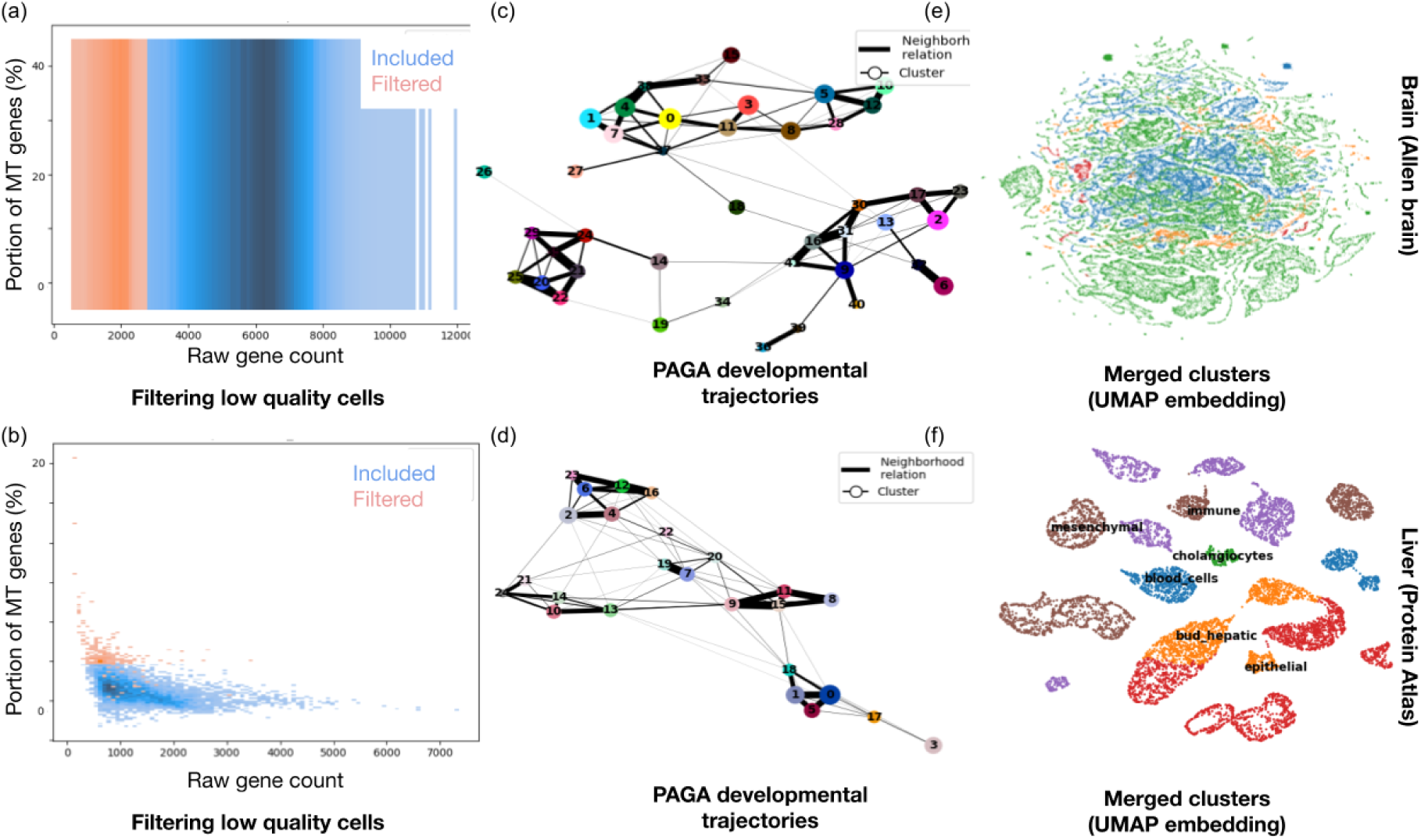
Single cells RNA-seq data prepossessing: cells with less than a threshold RNA-seq reads count or cells with a high portion of mitochondrial genes were removed (a,b). Cells with high-quality readds were then clustered on expression profile, and PAGA developmental trajectories were calculated (c, d). Obtained clusters were refined into merged sets of the same developmental stage and cell subtype(c, d).

To make sure all cells/samples have the same ratio of genes expressed for the deep learning model and other downstream analyses, sample counts for each cell were normalized by the total counts over all genes so that it sums up to 10 *×* 10^4^ and then pseudo-log-transformed these values as follows:

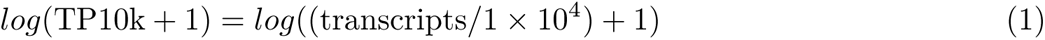

Datasets that combine microarray data^73^ (i.e the expression state of a large number of genes) are advantageous in achieving statistical power. However, we note that samples from different batches that were obtained and prepared at varying locations or times (e.g., GTEx) or comprise a large number of cells (e.g., Allen Brain Map) suffer from non-biological experimental variation or “batch effects”^74^. Batch effects often impose serious computational challenges and result in spurious outcomes and conclusions. Hence, to address this problem, we utilized DESC, an unsupervised deep embedding algorithm capable of removing batch effects that are smaller than the actual biological variation^56^. DESC also assigns cells with a more similar experimental setting into a soft cluster in an unsupervised manner^56^. For the DESC step, we adopted the default setting but only considered the maximum of 10 neighborhoods instead of 25 and incorporated densMAP community detection methodology for establishing seed clustering boundaries to better preserve the topological features of transcriptomic variability data^55, 75^.

GENIE3, GRNBoost2, PIDC and NS-DIMCORN account for cell sub-types and underlying cell state changes directly. For the rest of the algorithms that, in addition to RNA-seq data, require trajectory inference data as input, we inferred the progression of cells through geodesic distance along the coarse-grained map of the sequencing data manifold, based on the connectivity of manifold partitions [76] and provided these values to the algorithms as pseudo-times for mimicking real sequential cell states (c,d - Figures 6 and 7).

In order to avoid mischaracterizing sub-tissues as the “states of cells” by NS-DIMCORN, we further refined and merged the soft clusters previously assigned by DESC into larger sets dependent on consensus cell-type-specific marker genes of the brain and liver. Namely, brain cells were combined into five distinct groups of astrocytes, endothelial, microglia, neuron, and oligodendrocyte^77^; and liver cells were placed into either of the cholangiocytes, blood, mesenchymal, epithelial, immune and bud-hepatic sub-cluster^63^. Fine-grained final cell clusters (e,f - Figures 6 and 7) used to profile the robustness of NS-DIMCORN also accounted for developmental cell stages *G*_1_*/S*, *S*, *G*_2_ and *G*_2_*/M* , using well-characterized marker genes of essential cell cycle processes (DNA replication, chromosome segregation, and cell adhesion)^78^.

We included only the most relevant genetic drivers of each GRN, by identifying the set of most characteristic genes^45^ of each tissue/sample type and ranked them based on their variances across each single-cell thus increasing the power of downstream network inference algorithms. To further explain, after data standardization with a regularized standard deviation (i.e., z-score normalization per feature) and controlling for the relationship between mean expression and variability, the pseudo-log-transformed normalized variance of each gene was calculated as the variance of each gene; genes were then ranked by this variance^79^, and only the most highly variable 2500 genes were included in downstream analysis.

### 4.3 Estimating RNA-seq data real distribution

### 4.4 Estimating RNA-seq data real distribution

The main advantage of NS-DIMCORN is estimating continuous distributions of gene expression trajectories from RNA-seq data that allows sampling of the trajectory states between the available readouts as well as the observed results. We achieved the above by designing and optimizing an ODE network^80, 81^ that defines a continuous bijective map between vector field (latent variables, **y**) to RNA-seq data **x**, such that formally:

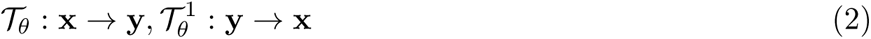

Sampling from high dimensional space of the RNA-seq data would be computationally expensive or infeasible, hence given the invertible function above, instead of directly parameterizing the distribution of RNA-seq data, we specified the data distribution implicitly by warping a base distribution **Z** *∼ p***_z_**(**z**), with an invertible (bijective) function. For RNA-seq data **x***_n,m_*with *m* genes, *n* observations and **x** *∈* **R***^D^* we chose **z**_0_ as the base distribution where **z**_0_ is a multivariate normal with *µ* = {*µ*_0_*, · · · , µ_m_*} and *σ* = {*σ*_0_*, · · · , σ_m_*}. Here *µ*_0_ is equal to the normalized mean of the first gene from *n* observations, and *σ*_0_ is its normalized variance among all the readouts for that gene. It should be noted that when the number of observations was less than 1280 (chosen based on the number of batches that could be fitted optimally on the available A100 Nvidia GPUs with 80 GB of memory) for a tissue/type of cell, in order to allow the model to converge with minimum fluctuation during the training, we augmented the dataset by generating new readouts within range of 0.1 ( just an arbitrary choice) of variance of each gene for randomly selected RNA-seq observations.

Thus, if the parameterized continuous dynamics of genes, trajectories using invertible ODE parametric function were specified by a neural network:

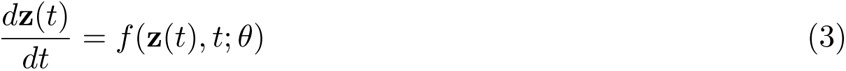

For training the network, NS-DIMCORN firstly takes samples from the base distribution **Z**_0_ *∼ p_z_*_0_ (**Z**_0_); then solves the initial value problem below:

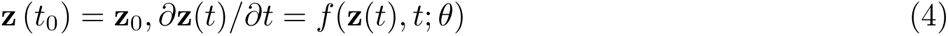

for **Z***_t_*_1_ using the Dormand-Prince explicit solver of non-stiff ODEs^82^ given the observations in the RNA-seq dataset. To calculate the value of **Z** (*t*_1_) *∈* **R***^D^*, the main challenge is computing the determinant of the Jacobian of the 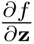, which can restrict the architecture of the neural network used^83^. Here we use an instantaneous change of variables Equation (5)^80^ as described by Chen et al. for this calculation that allows the gradients to be computed efficiently using the adjoint sensitivity method^80^:

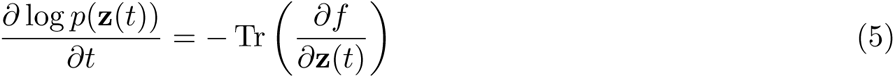

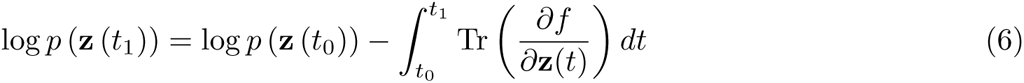

Also, to avoid the major pitfall of the RNA inference algorithms, instead of incorporating error-prone pseudo-time, we allowed the solver to choose *t_k−_*_1_ and *t_k_*, the time between two different observation, which was then integrated over time in Equation (6) as stated in Equation (5).

Eventually for every gene expression readout, NS-DIMCORN computed **Z**_0_ that generates that readout as well its likelihood using:

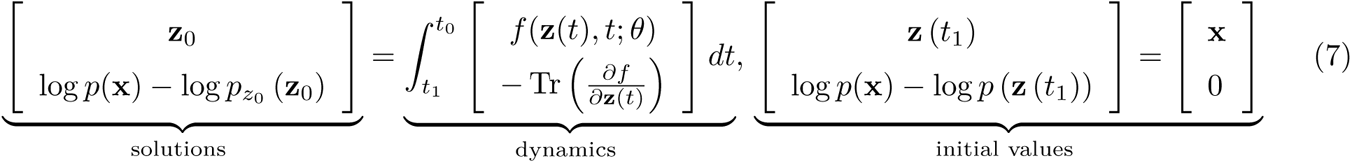

Given that now we can efficiently calculate the Jacobian of 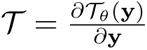 as follows we can keep track of the deformations using the change of variable formula (Equation (8)), and transfer the notion of probability onto **x** and invert it again if needed^83^ as follows:

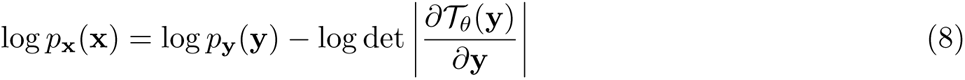

### 4.5 Co-variance Estimation

The inverse of the covariance matrix (precision matrix) is proportional to the partial independence relationship between matrix columns (genes). Under the assumption that only linear relationships exist between genes, if two genes are independent conditionally, all the corresponding coefficients in the precision matrix for those genes will be zero^84^. The covariance matrix of each sampled dataset can be calculated empirically. But inversion of the covariance matrix is computationally expensive and sometimes numerically impossible. Moreover, for high dimensional data or uncentered data samples, the precision matrix obtained from the inversion of the covariance matrix is not accurate (The Maximum Likelihood Estimator is not a good estimator of the eigenvalues of the covariance matrix). Consequently, estimating the precision matrix directly from data is the next best logical step^85^. To this end, we utilized a Hamiltonian Monte Carlo (HMC) sampler with adaptive step size^86^ where the target log probability was a multivariate normal, parameterized by Chelosky factors of the precision matrix and a Wishart distribution as prior distribution (conjugate prior of multivariate normal). The full implementation of the HMC sampler is described by MCMC using Hamiltonian dynamics paper^87^ and its implementation is described elsewhere^88^.

### 4.6 Mutual information (MI)

MI is a quantitative measurement of how much concurrent information exists about two variables. MI is a better measure of nonlinear interaction^59^ and consequently a good candidate for measuring non- linear interactions in GRNs. For two genes (*G_n_*, *G_m_* distributed according to some joint probability density *µ*(*g_n_, g_m_*), where marginal densities of *g_m_* is equal to *µ_g_* (*g_m_*) = ∫*dg_n_µ*(*g_m_, g_n_*) and the marginal densities of *g_n_* is equal to *µ_g_* (*g_n_*) = ∫ *dg_m_µ*(*g_n_, g_m_*) the MI is defined as

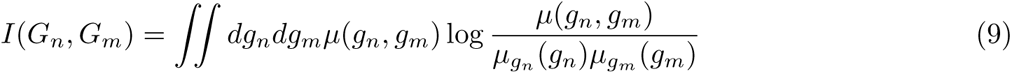

For a more efficient estimation of MI for strongly dependent variables (such are precursors of protein in a pathway), we tweaked the Kozachenko-Leonenko estimator^59^ for local nonuniformity correction such that if *V*(*i*) *⊂* **R***^d^* is the volume of *k* nearest neighbors of a sample point *g^i^* in some space; we assumed that there is some subset, 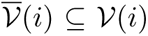 with volume *V* (*i*) *≤ V* (*i*) which density is constant as described by Geo et al., genes^60^ .

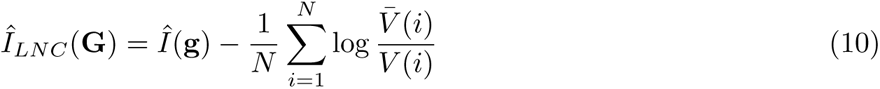

Thus, this correction term will improve the estimate of *V* (*i*) for strongly correlated interactions.

### 4.7 Model accuracy metrics for synthetic data

For synthetic data, where we were sure about the majority of the actual interactions (besides noise) in a GRE, we evaluated the result of each algorithm using the following criteria. We assigned the edges in the relevant network the true positive label and ranked edges from each method as the predictions. Beforehand, we omitted all self-loops given that some methods always assigned the highest rank to self-regulating genes where some other methods, such as SINGE, and critically NS-DIMCORN, ignore these interactions successfully.

#### 4.7.1 Precision (Pr)

The precision is the ratio

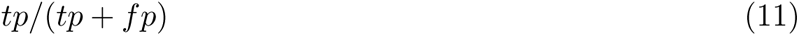

where tp is the number of true positives and fp is the number of false positives.

#### 4.7.2 The Area Under the Precision-Recall Curve (AUPRC)

AUPRC was calculated using the average_precision_ score function from sklearn^89^, which summarises the precision-recall curve as the weighted mean of precisions achieved at different thresholds.

#### 4.7.3 The Area Under the Receiver Operating Characteristic (AUROC)

AUROC was calculated using the roc_auc_score function from sklearn^89^. The Receiver Operating Characteristic (ROC) curve is created by plotting the true positive rate vs. the fraction of false positives out of the negative false positive rate at various threshold settings. A receiver operating characteristic curve, or ROC curve, illustrates the diagnostic ability of the classifier as its discrimination threshold changes. AUROC varies between 0 and 1, with 0.5 being an uninformative model.

#### 4.7.4 Balanced F-score (F1-S)

F1 scores compute the harmonic mean of precision and recall so that

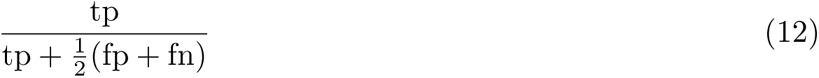

where tp is the number of true positives fn is false negatives and fp is the number of false positives. An F1 score of 1 is the best score and 0 is the worst possible F1 score.

#### 4.7.5 Correct Directions (CDiR)

CDiR is the portion of inferred interactions that also showed the direction of interaction between genes correctly in the biological sense this can be interpreted as inhibitory vs. excitatory regulation.

### 4.8 Model accuracy metrics for empirical data

Given that we could not fully establish which gene-gene interactions are false positives, true positives, and false negatives for empirical datasets, instead of the conventional accuracy metrics introduced above, we adopted a new set of modified accuracy measures for the bulk and the single-cell sequencing data. AUC and AUPRC estimates are sensitive to noisy data^90^ yet are linearly related to observed accuracy^91^ and it is also proven that both are closely related to the Wilcoxon test of ranks^92, 93^. Using the above line of reasoning, we assumed that we could extrapolate the accuracy of a model from a portion of identified true positives and false negatives. Hence for predictor model *f* , we defined True in the Top 5000 Identified Interactions (T5KI) as an unbiased surrogate metric of the model’s discrimination and calibration. T5KI can be interpreted like the Early Precision Ratio (EPR) metric that was previously used for benchmarking GRN inference algorithms^8^.

#### 4.8.1 All Interactions Identified (AII)

AII counts the number of all the identified interactions.

#### 4.8.2 Genuine Identified Interactions (GII)

In the context of empirical data, AII measures the number of identified interactions identical to those in our curated super pathway dataset.

#### 4.8.3 Missing Expected Interactions (MEI)

MEI is the offset of unidentified interactions from the supers pathway dataset.

#### 4.8.4 Genuine Interaction in the Top 5000 Identified Interactions (T5KI)

T5KI measures the number of AII only in the top-ranked 5000 inferred interactions while incurring the penalty for missing known-to-be true interactions (MEI). Formally

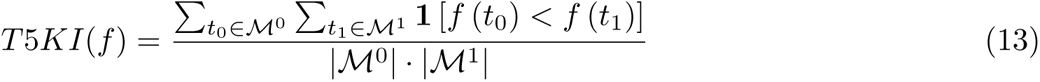

here **1** [*f* (*t*_0_) *< f* (*t*_1_)] denotes an indicator function which returns 1 iff *f* (*t*_0_) *< f* (*t*_1_) otherwise return 0; *ℳ*^0^ is the set of negative examples such as MEI, and *M*^1^ is the set of positive examples such as AII.

### 4.9 Graph topology metrics

For the strictly defined weighted Graph *G* with sets of vertices *G* ⊆ {*v*_1_*, …, v_n_*}, *n* nodes and *m* edges

#### 4.9.1 Graph Path Length (GPL)

is the normalized sum of path lengths *d*(*s, t*) between all pairs of nodes, and it measures the efficiency of information flow for a network. Here we reported the averaged PL of each sub-graph when disconnected graphs were observed.

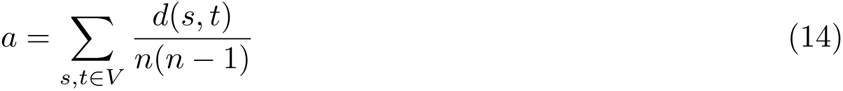

#### 4.9.2 The Averaged Degree (D)

is the averaged number of adjacent edges to nodes, while Averaged Degree Centrality (DC) is the portion of nodes connected to each node. The Density (Den) of network *G* is defined as

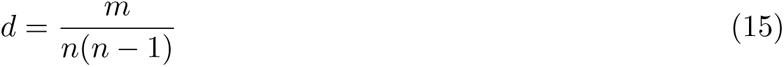

Together, *D, DC*, and *Den* were used as a surrogate for the first moment and second moment of the Degree distribution, which indicates the number and size of the network hubs.

#### 4.9.3 Graph Modularity (GM)

uses Clauset*−*Newman*−*Moore greedy modularity maximization and calculates the strength of divisions of the network into clusters.

#### 4.9.4 Graph clustering Coefficient (GCE)

has been calculated as the geometric average of the sub-graphs normalized edge weights and measures the degree to which nodes in a graph tend to cluster together. Formally

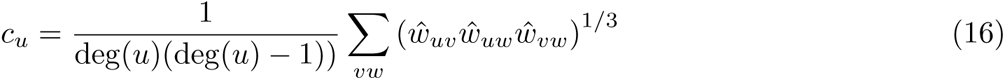

Where 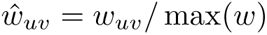

#### 4.9.5 Graph Diameter (GDm)

is the measure of the graph’s eccentricity. In other words, the maximum distance between a vertex to all other vertices is called the diameter, Dm. Dm is in contrast to the more behavioral metrics discussed beforehand (how each node behaves), *M, Ce*, and *Dm* focus on the topological level of a network.

#### 4.9.6 Graph Distances (GDi)

We calculated structural distance *D* (Γ_1_, Γ_2_) between two different graphs in terms of the Jensen-Shannon distance J S of Laplacian spectra of the two graphs^94^. Briefly if Gaussian kernel 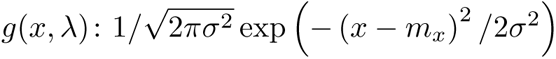 exists the function of convoluted spectrum of a network with *σ* = .01 is defined as

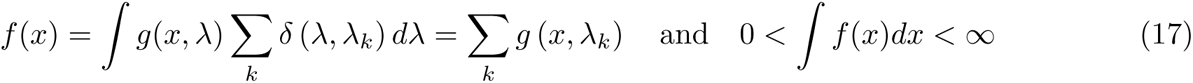

spectral density *f ^∗^* was then calculated by normalizing *f* as:

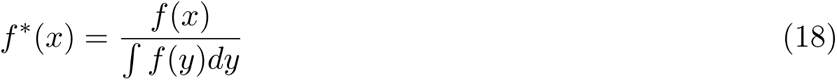

and distance is equal to

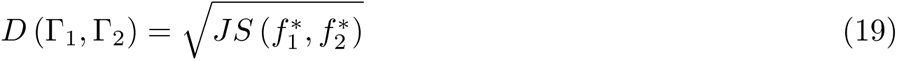

#### 4.9.7 Graph Fidelity (GFid)

Fi combines the properties of a complex inferred graph and ground truth networks to compare their sim- ilarity overall with a single numerical calculate fidelity metric *δ* as described by Alexandru Topirceanu^37^. Fidelity measures the averages over an arbitrary number of measurements for a graph.

### 4.10 Overview of the benchmarked algorithms

#### 4.10.1 GENIE3

GENIE3^25^ was the top performer algorithm for inferring regulatory networks for bulk transcriptional data in the DREAM4 challenge. GENIE3 regresses the expression profile of genes one at a time and then ranks each gene’s importance in predicting other genes’ profiles using random forests. It then constructs regulatory networks by aggregating these weights such that the level of importance becomes the edge weights in the network.

#### 4.10.2 GRISLI

GRISLI^30^ uses linear ODEs to calculate how gene expression values change during the cell sampled cell states for the provided experimental times (here, input pseudo-times).

#### 4.10.3 GRNBoost2

GRNBoost2^31^ uses regression and tree-based models like GENIE3 but incorporates stochastic gradient boosting and early stopping to achieve better speed for bigger networks with more genes under study.

#### 4.10.4 LEAP

LEAP^17^ calculates asymmetric Pearson correlation of RNA-seq read counts between permuted different experimental times. The Maximum Pearson’s correlation between two pairs indicates the directed edge weights in the network.

#### 4.10.5 PIDC

PIDC^18^ calculates unique mutual information between two genes such that the relationship between two genes is proportional to the relationship of those genes to all the other genes in the network.

#### 4.10.6 PPCOR

PPCOR^15^ estimates the pairwise partial correlation coefficients given all the other genes’ expressions and computes a P-value for each correlation. Negative correlations here are deemed inhibitory and positive correlations are considered activating.

#### 4.10.7 SCODE

SCODE^32^ is essentially the data dimensionality and regress for linear ODEs to describe how gene-gene interactions result in observed gene expression dynamics.

#### 4.10.8 SCRIBE

SCRIBE^33^ computes mutual information between the past state of a regulator gene and the current state of a regulated gene, given the state of the regulated gene at the last experimental time, then excludes interactions relating to other indirect effects.

#### 4.10.9 SINCERITIES

SINCERITIES^19^ infers gene-gene weights in GRNs by computing changes in the distributions of gene ex- pressions between two consecutive experimental times using the Kolmogorov–Smirnov statistic and ridge regression. Partial correlation analyses between pairs of genes then indicate the direction interactions in these networks.

## 5 Supplementary information

**Table S1.**
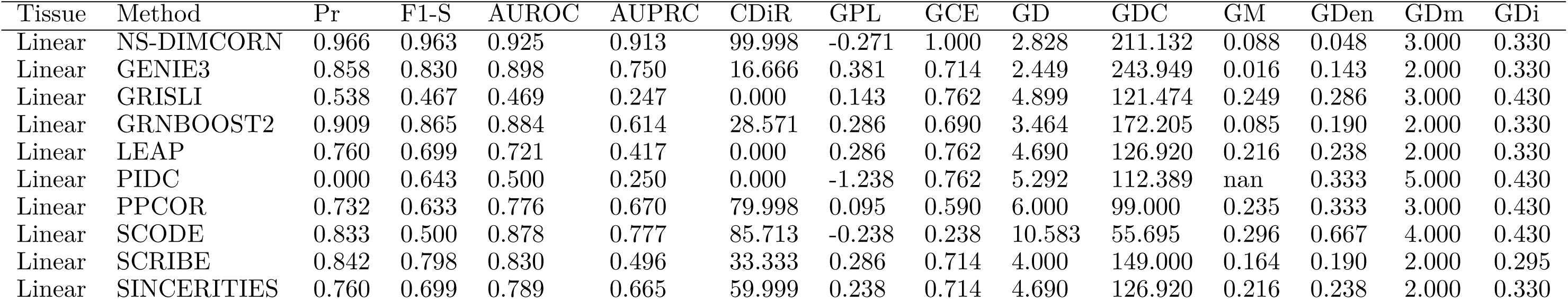
Benchmarking NS-DIMCORN against the other state-of-art GRN inference algorithms for synthetic data that represents linear trajectory for cellular process as depicted at fig. 2

| Tissue | Method | Pr | F1-S | AUROC | AUPRC | CDiR | GPL | GCE | GD | GDC | GM | GDen | GDm | GDi |
| --- | --- | --- | --- | --- | --- | --- | --- | --- | --- | --- | --- | --- | --- | --- |
| Linear | NS-DIMCORN | 0.966 | 0.963 | 0.925 | 0.913 | 99.998 | -0.271 | 1.000 | 2.828 | 211.132 | 0.088 | 0.048 | 3.000 | 0.330 |
| Linear | GENIE3 | 0.858 | 0.830 | 0.898 | 0.750 | 16.666 | 0.381 | 0.714 | 2.449 | 243.949 | 0.016 | 0.143 | 2.000 | 0.330 |
| Linear | GRISLI | 0.538 | 0.467 | 0.469 | 0.247 | 0.000 | 0.143 | 0.762 | 4.899 | 121.474 | 0.249 | 0.286 | 3.000 | 0.430 |
| Linear | GRNBOOST2 | 0.909 | 0.865 | 0.884 | 0.614 | 28.571 | 0.286 | 0.690 | 3.464 | 172.205 | 0.085 | 0.190 | 2.000 | 0.330 |
| Linear | LEAP | 0.760 | 0.699 | 0.721 | 0.417 | 0.000 | 0.286 | 0.762 | 4.690 | 126.920 | 0.216 | 0.238 | 2.000 | 0.330 |
| Linear | PIDC | 0.000 | 0.643 | 0.500 | 0.250 | 0.000 | -1.238 | 0.762 | 5.292 | 112.389 | nan | 0.333 | 5.000 | 0.430 |
| Linear | PPCOR | 0.732 | 0.633 | 0.776 | 0.670 | 79.998 | 0.095 | 0.590 | 6.000 | 99.000 | 0.235 | 0.333 | 3.000 | 0.430 |
| Linear | SCODE | 0.833 | 0.500 | 0.878 | 0.777 | 85.713 | -0.238 | 0.238 | 10.583 | 55.695 | 0.296 | 0.667 | 4.000 | 0.430 |
| Linear | SCRIBE | 0.842 | 0.798 | 0.830 | 0.496 | 33.333 | 0.286 | 0.714 | 4.000 | 149.000 | 0.164 | 0.190 | 2.000 | 0.295 |
| Linear | SINCERITIES | 0.760 | 0.699 | 0.789 | 0.665 | 59.999 | 0.238 | 0.714 | 4.690 | 126.920 | 0.216 | 0.238 | 2.000 | 0.330 |

**Table S2.**
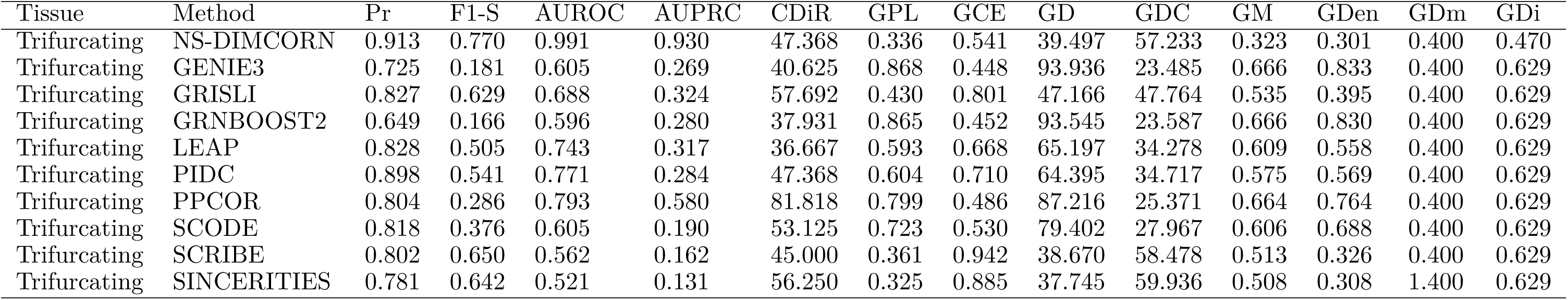
Benchmarking NS-DIMCORN against the other state-of-art GRN inference algorithms for synthetic data that represents trifurcating trajectory for cellular process as depicted at fig. 2

**Table S3.** Benchmarking NS-DIMCORN against the other state-of-art GRN inference algorithms for synthetic data that represents cyclic trajectory for cellular process as depicted at fig. 2

| Tissue | Method | Pr | F1-S | AUROC | AUPRC | CDiR | GPL | GCE | GD | GDC | GM | GDen | GDm | GDi |
| --- | --- | --- | --- | --- | --- | --- | --- | --- | --- | --- | --- | --- | --- | --- |
| Cyclic | NS-DIMCORN | 0.838 | 0.816 | 0.944 | 0.883 | 99.998 | 0.800 | 0.417 | 3.162 | 157.114 | 0.049 | 0.133 | 0.000 | 0.284 |
| Cyclic | GENIE3 | 0.714 | 0.714 | 0.722 | 0.687 | 33.332 | 0.350 | 0.444 | 2.000 | 249.000 | 0.111 | 1.000 | 1.500 | 0.572 |
| Cyclic | GRISLI | 0.813 | 0.772 | 0.789 | 0.565 | 20.000 | 0.800 | 0.389 | 3.464 | 143.338 | 0.043 | 0.200 | 0.000 | 0.314 |
| Cyclic | GRNBOOST2 | 0.752 | 0.755 | 0.700 | 0.677 | 33.332 | 0.433 | 0.611 | 2.000 | 249.000 | 0.153 | 0.067 | 1.500 | 0.464 |
| Cyclic | LEAP | 0.752 | 0.755 | 0.856 | 0.771 | 33.332 | 0.433 | 0.611 | 2.000 | 249.000 | 0.153 | 0.067 | 1.500 | 0.464 |
| Cyclic | PIDC | 0.959 | 0.953 | 0.967 | 0.915 | 49.999 | 0.867 | 0.750 | 2.449 | 203.124 | 0.044 | 0.067 | 0.000 | 0.284 |
| Cyclic | PPCOR | 0.929 | 0.908 | 0.911 | 0.697 | 99.998 | 0.733 | 0.500 | 2.449 | 203.124 | 0.049 | 0.133 | 1.000 | 0.579 |
| Cyclic | SCODE | 0.754 | 0.635 | 0.789 | 0.645 | 39.999 | 0.467 | 0.156 | 5.292 | 93.491 | 0.167 | 0.400 | 1.000 | 0.395 |
| Cyclic | SCRIBE | 0.714 | 0.714 | 0.844 | 0.729 | 66.664 | 1.000 | 1.000 | 1.414 | 352.553 | 0.000 | 1.000 | 0.000 | nan |
| Cyclic | SINCERITIES | 0.619 | 0.619 | 0.544 | 0.306 | 99.995 | -0.033 | 0.500 | 3.162 | 157.114 | 0.167 | 1.000 | 2.000 | 0.464 |

**Table S4.** Benchmarking NS-DIMCORN against the other state-of-art GRN inference algorithms for synthetic data that represents compound trajectories for cellular processes as depicted at fig. 2

| Tissue | Method | Pr | F1-S | AUROC | AUPRC | CDiR | GPL | GCE | GD | GDC | GM | GDen | GDm | GDi |
| --- | --- | --- | --- | --- | --- | --- | --- | --- | --- | --- | --- | --- | --- | --- |
| Compound | NS-DIMCORN | 0.939 | 0.350 | 0.997 | 0.925 | 61.224 | 0.352 | 0.280 | 180.159 | 19.537 | 0.719 | 0.765 | 1.200 | 0.395 |
| Compound | GENIE3 | 0.892 | 0.192 | 0.726 | 0.217 | 60.000 | 0.247 | 0.257 | 199.369 | 17.559 | 0.765 | 0.871 | 1.200 | 0.395 |
| Compound | GRISLI | 0.903 | 0.462 | 0.697 | 0.226 | 62.500 | 0.464 | 0.382 | 155.603 | 22.778 | 0.718 | 0.653 | 1.200 | 0.395 |
| Compound | GRNBOOST2 | 0.893 | 0.155 | 0.759 | 0.228 | 58.696 | 0.221 | 0.239 | 204.828 | 17.064 | 0.779 | 0.896 | 1.200 | 0.395 |
| Compound | LEAP | 0.918 | 0.505 | 0.766 | 0.210 | 53.488 | 0.499 | 0.435 | 146.219 | 24.305 | 0.721 | 0.619 | 1.200 | 0.395 |
| Compound | PIDC | 0.932 | 0.661 | 0.836 | 0.214 | 60.000 | 0.664 | 0.643 | 106.558 | 33.723 | 0.660 | 0.454 | 1.200 | 0.395 |
| Compound | PPCOR | 0.934 | 0.362 | 0.899 | 0.676 | 85.416 | 0.363 | 0.299 | 176.969 | 19.908 | 0.751 | 0.754 | 1.200 | 0.395 |
| Compound | SCODE | 0.938 | 0.106 | 0.542 | 0.132 | 61.224 | 0.187 | 0.211 | 212.186 | 16.438 | 0.779 | 0.930 | 2.200 | 0.395 |
| Compound | SCRIBE | 0.889 | 0.651 | 0.573 | 0.089 | 57.143 | 0.693 | 0.708 | 98.271 | 36.651 | 0.661 | 0.424 | 1.200 | 0.395 |
| Compound | SINCERITIES | 0.886 | 0.662 | 0.565 | 0.085 | 57.692 | 0.712 | 0.675 | 96.287 | 37.427 | 0.642 | 0.405 | 1.200 | 0.395 |

**Table S5.**
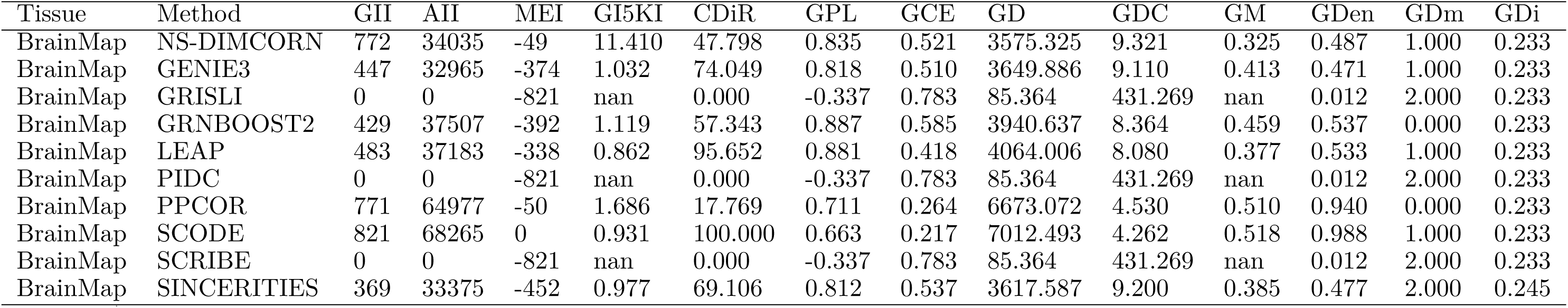
Benchmarking NS-DIMCORN against the other state-of-art GRN inference algorithms using brain scRNA-seq data obtained from BrainMap fig. 5

**Table S6.** Benchmarking NS-DIMCORN against the other state-of-art GRN inference algorithms using brain scRNA-seq data obtained from Protein Atlas fig. 5

| Tissue | Method | GII | AII | MEI | GI5KI | CDiR | GPL | GCE | GD | GDC | GM | GDen | GDm | GDi |
| --- | --- | --- | --- | --- | --- | --- | --- | --- | --- | --- | --- | --- | --- | --- |
| LiverMap | NS-DIMCORN | 1384 | 46983 | -53 | 10.008 | 55.058 | 0.797 | 0.528 | 4833.467 | 7.958 | 0.457 | 0.485 | 0.111 | 0.225 |
| LiverMap | GENIE3 | 918 | 49412 | -519 | 2.433 | 77.233 | 0.822 | 0.611 | 4831.999 | 7.961 | 0.419 | 0.511 | 1.111 | 0.225 |
| LiverMap | GRISLI | 1032 | 64163 | -405 | 1.741 | 100.000 | -0.031 | 0.387 | 6498.063 | 5.664 | 0.486 | 0.668 | 1.089 | 0.469 |
| LiverMap | GRNBOOST2 | 974 | 60753 | -463 | 3.219 | 61.499 | 0.943 | 0.597 | 5784.176 | 6.486 | 0.472 | 0.631 | 0.111 | 0.225 |
| LiverMap | LEAP | 901 | 45906 | -536 | 2.629 | 96.892 | 0.777 | 0.502 | 4549.114 | 8.518 | 0.293 | 0.473 | 1.111 | 0.225 |
| LiverMap | PIDC | 894 | 46708 | -543 | 3.051 | 84.116 | 0.794 | 0.639 | 4499.431 | 8.623 | 0.359 | 0.482 | 1.111 | 0.225 |
| LiverMap | PPCOR | 1358 | 90160 | -79 | 3.160 | 14.948 | 0.744 | 0.325 | 8520.335 | 4.082 | 0.509 | 0.944 | 0.111 | 0.225 |
| LiverMap | SCODE | 1437 | 93961 | 0 | 1.935 | 100.000 | 0.703 | 0.285 | 8882.588 | 3.875 | 0.514 | 0.985 | 0.889 | 0.225 |
| LiverMap | SINCERITIES | 904 | 52193 | -533 | 2.817 | 68.363 | 0.852 | 0.604 | 5007.700 | 7.647 | 0.367 | 0.540 | 0.111 | 0.225 |

**Table S7.** Benchmarking NS-DIMCORN against the other state-of-art GRN inference algorithms using brain RNA-seq data obtained from GTEx fig. 5

| Tissue | Method | GII | AII | MEI | GI5KI | CDiR | GPL | GCE | GD | GDC | GM | GDen | GDm | GDi |
| --- | --- | --- | --- | --- | --- | --- | --- | --- | --- | --- | --- | --- | --- | --- |
| GTEXBrain | NS-DIMCORN | 617 | 33700 | -112 | 9.810 | 59.643 | 0.771 | 0.658 | 3529.475 | 9.341 | 0.524 | 0.494 | 0.167 | 0.237 |
| GTEXBrain | GENIE3 | 552 | 49199 | -177 | 3.280 | 44.203 | 0.997 | 0.451 | 5167.502 | 6.063 | 0.589 | 0.726 | 0.167 | 0.237 |
| GTEXBrain | GRISLI | 647 | 57676 | -82 | 1.498 | 100.000 | 0.300 | 0.286 | 6146.067 | 4.939 | 0.618 | 0.853 | 0.333 | 0.560 |
| GTEXBrain | GRNBOOST2 | 636 | 58207 | -93 | 3.222 | 35.220 | 0.862 | 0.361 | 6039.692 | 5.043 | 0.613 | 0.861 | 0.167 | 0.237 |
| GTEXBrain | LEAP | 418 | 32146 | -311 | 2.215 | 94.498 | 0.739 | 0.542 | 3465.000 | 9.534 | 0.437 | 0.470 | 1.167 | 0.237 |
| GTEXBrain | PIDC | 380 | 32989 | -349 | 1.840 | 72.105 | 0.760 | 0.581 | 3526.582 | 9.350 | 0.465 | 0.483 | 1.167 | 0.237 |
| GTEXBrain | PPCOR | 484 | 39033 | -245 | 3.290 | 36.570 | 0.851 | 0.640 | 4050.023 | 8.012 | 0.593 | 0.573 | 0.167 | 0.237 |
| GTEXBrain | SCODE | 729 | 66795 | 0 | 2.456 | 100.000 | 0.733 | 0.249 | 6906.650 | 4.285 | 0.628 | 0.989 | 0.833 | 0.237 |
| GTEXBrain | SCRIBE | 0 | 0 | -729 | nan | 0.000 | -0.267 | 0.751 | 76.211 | 477.934 | nan | 0.011 | 1.833 | 0.237 |
| GTEXBrain | SINCERITIES | 395 | 34760 | -334 | 1.302 | 83.038 | 0.787 | 0.655 | 3640.856 | 9.025 | 0.525 | 0.509 | 0.167 | 0.237 |

**Table S8.** Benchmarking NS-DIMCORN against the other state-of-art GRN inference algorithms using liver RNA-seq data obtained from GTEx fig. 5

| Tissue | Method | GII | AII | MEI | GI5KI | CDiR | GPL | GCE | GD | GDC | GM | GDen | GDm | GDi |
| --- | --- | --- | --- | --- | --- | --- | --- | --- | --- | --- | --- | --- | --- | --- |
| GTExLiver | NS-DIMCORN | 1109 | 60832 | -133 | 14.679 | 52.119 | 0.939 | 0.210 | 5449.034 | 8.047 | 0.388 | 0.489 | 0.714 | 0.210 |
| GTExLiver | GENIE3 | 263 | 15753 | -979 | 1.581 | 100.000 | -0.558 | 0.861 | 2296.129 | 20.471 | 0.512 | 0.119 | 2.711 | 0.210 |
| GTExLiver | GRISLI | 1199 | 116005 | -43 | 1.509 | 100.000 | -0.040 | 0.231 | 10436.040 | 3.724 | 0.511 | 0.942 | 1.214 | 0.258 |
| GTExLiver | GRNBOOST2 | 1000 | 96794 | -242 | 3.794 | 36.000 | 0.644 | 0.403 | 8663.514 | 4.691 | 0.495 | 0.785 | 0.714 | 0.210 |
| GTExLiver | LEAP | 635 | 55230 | -607 | 2.018 | 88.189 | 0.986 | 0.582 | 5086.391 | 8.693 | 0.337 | 0.443 | 0.286 | 0.210 |
| GTExLiver | PIDC | 0 | 0 | -1242 | nan | 0.000 | -0.561 | 0.779 | 111.760 | 440.122 | nan | 0.010 | 2.714 | 0.210 |
| GTExLiver | PPCOR | 1242 | 121771 | 0 | 0.000 | 0.000 | 0.439 | 0.221 | 10845.712 | 3.546 | 0.512 | 0.990 | 1.714 | 0.210 |
| GTExLiver | SCODE | 1242 | 121771 | 0 | 1.216 | 100.000 | 0.439 | 0.221 | 10845.712 | 3.546 | 0.512 | 0.990 | 1.714 | 0.210 |
| GTExLiver | SCRIBE | 790 | 73887 | -452 | 2.394 | 86.709 | 0.832 | 0.551 | 6678.529 | 6.382 | 0.423 | 0.597 | 0.714 | 0.210 |
| GTExLiver | SINCERITIES | 747 | 70082 | -495 | 1.013 | 99.331 | 1.000 | 0.472 | 6486.310 | 6.601 | 0.401 | 0.565 | 1.214 | 0.210 |

## Notes

### Competing Interest Statement

The authors have declared no competing interest.

## References

1. Stark, R., Grzelak, M. & Hadfield, J. RNA sequencing: the teenage years. en. Nat. Rev. Genet. 20, 631–656 (Nov. 2019).

2. Aalto, A., Viitasaari, L., Ilmonen, P., Mombaerts, L. & Gonçalves, J. Gene regulatory network inference from sparsely sampled noisy data. en. Nat. Commun. 11, 3493 (July 2020).

3. Yu, L., Fernandez, S. & Brock, G. Power analysis for RNA-Seq differential expression studies. en. BMC Bioinformatics 18, 234 (May 2017).

4. Zhao, M., He, W., Tang, J., Zou, Q. & Guo, F. A comprehensive overview and critical evaluation of gene regulatory network inference technologies. en. Brief. Bioinform. 22 (Sept. 2021).

5. Singh, D., Singh, P. K., Chaudhary, S., Mehla, K. & Kumar, S. in Advances in Genetics (eds Friedmann, T., Dunlap, J. C. & Goodwin, S. F.) 87–121 (Academic Press, Jan. 2012).

6. Cardoso, T. F., et al. RNA-seq based detection of differentially expressed genes in the skeletal muscle of Duroc pigs with distinct lipid profiles. en. Sci. Rep. 7, 40005 (Feb. 2017).

7. Statello, L., Guo, C.-J., Chen, L.-L. & Huarte, M. Gene regulation by long non-coding RNAs and its biological functions. en. Nat. Rev. Mol. Cell Biol. 22, 96–118 (Feb. 2021).

8. Pratapa, A., Jalihal, A. P., Law, J. N., Bharadwaj, A. & Murali, T. M. Benchmarking algorithms for gene regulatory network inference from single-cell transcriptomic data. en. Nat. Methods 17, 147–154 (Feb. 2020).

9. Alberch, P. Kauffman, S. A. The origins of order. Self-organization and selection in evolution. Oxford University Press (1993). Price: f17.95 (pb), f51.00 (hb). ISBN: 0-19-505811-9 (hb) and 0-19-507951-5 (pb). en. J. Evol. Biol. 7, 518–519 (July 1994).

10. Borriello, E. & Daniels, B. C. The basis of easy controllability in Boolean networks. en. Nat. Commun. 12, 5227 (Sept. 2021).

11. Huynh-Thu, V. A., Irrthum, A., Wehenkel, L. & Geurts, P. Inferring regulatory networks from expression data using tree-based methods. en. PLoS One 5 (Sept. 2010).

12. Woodhouse, S., Piterman, N., Wintersteiger, C. M., Göttgens, B. & Fisher, J. SCNS: a graphical tool for reconstructing executable regulatory networks from single-cell genomic data. en. BMC Syst. Biol. 12, 59 (May 2018).

13. Omranian, N., Eloundou-Mbebi, J. M. O., Mueller-Roeber, B. & Nikoloski, Z. Gene regulatory network inference using fused LASSO on multiple data sets. en. Sci. Rep. 6, 20533 (Feb. 2016).

14. Sanchez-Castillo, M., Blanco, D., Tienda-Luna, I. M., Carrion, M. C. & Huang, Y. A Bayesian framework for the inference of gene regulatory networks from time and pseudo-time series data. en. Bioinformatics 34, 964–970 (Mar. 2018).

15. Kim, S. Ppcor: An R package for a fast calculation to semi-partial correlation coefficients. en. Commun. Stat. Appl. Methods 22, 665–674 (Nov. 2015).

16. Opgen-Rhein, R. & Strimmer, K. From correlation to causation networks: a simple approximate learning algorithm and its application to high-dimensional plant gene expression data. en. BMC Syst. Biol. 1, 37 (Aug. 2007).

17. Specht, A. T. & Li, J. LEAP: constructing gene co-expression networks for single-cell RNA-sequencing data using pseudotime ordering. en. Bioinformatics 33, 764–766 (Mar. 2017).

18. Chan, T. E., Stumpf, M. P. H. & Babtie, A. C. Gene Regulatory Network Inference from Single-Cell Data Using Multivariate Information Measures. en. Cell Syst 5, 251–267.e3 (Sept. 2017).

19. Papili Gao, N., Ud-Dean, S. M. M., Gandrillon, O. & Gunawan, R. SINCERITIES: inferring gene regulatory networks from time-stamped single cell transcriptional expression profiles. en. Bioinformatics 34, 258–266 (Jan. 2018).

20. Zhao, H. & Duan, Z.-H. Cancer Genetic Network Inference Using Gaussian Graphical Models. en. Bioinform. Biol. Insights 13, 1177932219839402 (Apr. 2019).

21. Bennasar, M., Hicks, Y. & Setchi, R. Feature selection using Joint Mutual Information Maximisation. Expert Syst. Appl. 42, 8520–8532 (Dec. 2015).

22. Aljabbouli, H., Albizri, A. & Harfouche, A. Tree-Based Algorithm for Stable and Efficient Data Clustering. en. Informatics 7, 38 (Sept. 2020).

23. Aderhold, A., Husmeier, D. & Grzegorczyk, M. Approximate Bayesian inference in semi-mechanistic models. en. Stat. Comput. 27, 1003–1040 (2017).

24. Casadiego, J., Nitzan, M., Hallerberg, S. & Timme, M. Model-free inference of direct network interactions from nonlinear collective dynamics. en. Nat. Commun. 8, 2192 (Dec. 2017).

25. Huynh-Thu, V. A. & Geurts, P. dynGENIE3: dynamical GENIE3 for the inference of gene networks from time series expression data. en. Sci. Rep. 8, 3384 (Feb. 2018).

26. Mangan, N. M., Brunton, S. L., Proctor, J. L. & Kutz, J. N. Inferring Biological Networks by Sparse Identification of Nonlinear Dynamics. *IEEE Transactions on Molecular*, Biological and Multi-Scale Communications 2, 52–63 (June 2016).

27. Bansal, M., Della Gatta, G. & di Bernardo, D. Inference of gene regulatory networks and compound mode of action from time course gene expression profiles. en. Bioinformatics 22, 815–822 (Apr. 2006).

28. Penfold, C. A., Shifaz, A., Brown, P. E., Nicholson, A. & Wild, D. L. CSI: a nonparametric Bayesian approach to network inference from multiple perturbed time series gene expression data. en. Stat. Appl. Genet. Mol. Biol. 14, 307–310 (June 2015).

29. Paninski, L. Estimation of entropy and mutual information. en. Neural Comput. 15, 1191–1253 (June 2003).

30. Aubin-Frankowski, P.-C. & Vert, J.-P. Gene regulation inference from single-cell RNA-seq data with linear differential equations and velocity inference. en. Bioinformatics 36, 4774–4780 (Sept. 2020).

31. Moerman, T., et al. GRNBoost2 and Arboreto: efficient and scalable inference of gene regulatory networks. en. Bioinformatics 35, 2159–2161 (June 2019).

32. Matsumoto, H., et al. SCODE: an efficient regulatory network inference algorithm from single-cell RNA-Seq during differentiation. en. Bioinformatics 33, 2314–2321 (Aug. 2017).

33. Qiu, X., et al. Inferring Causal Gene Regulatory Networks from Coupled Single-Cell Expression Dynamics Using Scribe. en. Cell Syst 10, 265–274.e11 (Mar. 2020).

34. Irwin, M. & Wang, Z. Dynamic Systems Modeling Aug. 2017.

35. Bahadorian, M., et al. A topology-dynamics-based control strategy for multi-dimensional complex networked dynamical systems. en. Sci. Rep. 9, 19831 (Dec. 2019).

36. Sutherland, W. A. Introduction to Metric and Topological Spaces (Oxford Mathematics) 2nd ed. en (Oxford University Press, Oct. 2009).

37. Topirceanu, A., Udrescu, M. & Vladutiu, M. Network Fidelity: A Metric to Quantify the Similarity and Realism of Complex Networks in 2013 International Conference on Cloud and Green Computing (Sept. 2013), 289–296.

38. Preciado, V. M., Jadbabaie, A. & Verghese, G. C. Structural Analysis of Laplacian Spectral Properties of Large-Scale Networks. IEEE Trans. Automat. Contr. 58, 2338–2343 (Sept. 2013).

39. Lozoya, O. A., Santos, J. H. & Woychik, R. P. A Leveraged Signal-to-Noise Ratio (LSTNR) Method to Extract Differentially Expressed Genes and Multivariate Patterns of Expression From Noisy and Low- Replication RNAseq Data. en. Front. Genet. 9, 176 (May 2018).

40. Saelens, W., Cannoodt, R., Todorov, H. & Saeys, Y. A comparison of single-cell trajectory inference meth- ods. en. Nat. Biotechnol. 37, 547–554 (May 2019).

41. Gilpin, L. H., et al. Explaining Explanations: An Overview of Interpretability of Machine Learning in 2018 IEEE 5th International Conference on Data Science and Advanced Analytics (DSAA) (Oct. 2018), 80–89.

42. *BoolODE — BEELINE documentation* en. https://murali-group.github.io/Beeline/BoolODE.html. Accessed: 2023-4-14.

43. Hong, M., et al. RNA sequencing: new technologies and applications in cancer research. en. J. Hematol. Oncol. 13, 166 (Dec. 2020).

44. Datlinger, P., et al. Ultra-high-throughput single-cell RNA sequencing and perturbation screening with combinatorial fluidic indexing. en. Nat. Methods 18, 635–642 (June 2021).

45. Harrow, J., et al. GENCODE: the reference human genome annotation for The ENCODE Project. en. Genome Res. 22, 1760–1774 (Sept. 2012).

46. Horlbeck, M. A., et al. Mapping the Genetic Landscape of Human Cells. en. Cell 174, 953–967.e22 (Aug. 2018).

47. Li, X. & Wang, C.-Y. From bulk, single-cell to spatial RNA sequencing. en. Int. J. Oral Sci. 13, 36 (Nov. 2021).

48. Kharchenko, P. V. The triumphs and limitations of computational methods for scRNA-seq. en. Nat. Methods 18, 723–732 (July 2021).

49. Szklarczyk, D., et al. The STRING database in 2021: customizable protein-protein networks, and functional characterization of user-uploaded gene/measurement sets. en. Nucleic Acids Res. 49, D605–D612 (Jan. 2021).

50. Subramanian, A., et al. Gene set enrichment analysis: a knowledge-based approach for interpreting genome- wide expression profiles. en. Proc. Natl. Acad. Sci. U. S. A. 102, 15545–15550 (Oct. 2005).

51. Mubeen, S., et al. The Impact of Pathway Database Choice on Statistical Enrichment Analysis and Predictive Modeling. en. Front. Genet. 10, 1203 (Nov. 2019).

52. Deshpande, A., Chu, L.-F., Stewart, R. & Gitter, A. Network inference with Granger causality ensembles on single-cell transcriptomics. en. Cell Rep. 38, 110333 (Feb. 2022).

53. Abadi, M. et al. TensorFlow: Large-Scale Machine Learning on Heterogeneous Distributed Systems.

54. cupy: NumPy & SciPy for GPU en.

55. Narayan, A., Berger, B. & Cho, H. Assessing single-cell transcriptomic variability through density-preserving data visualization. en. Nat. Biotechnol. (Jan. 2021).

56. Li, X., et al. Deep learning enables accurate clustering with batch effect removal in single-cell RNA-seq analysis. en. Nat. Commun. 11, 2338 (May 2020).

57. McInnes, L., Healy, J. & Melville, J. UMAP: Uniform Manifold Approximation and Projection for Dimension Reduction. arXiv: 1802.03426 [stat.ML] (Feb. 2018).

58. Song, L., Langfelder, P. & Horvath, S. Comparison of co-expression measures: mutual information, correlation, and model based indices. en. BMC Bioinformatics 13, 328 (Dec. 2012).

59. Kraskov, A., Stögbauer, H. & Grassberger, P. Estimating mutual information. en. Phys. Rev. E Stat. Nonlin. Soft Matter Phys. 69, 066138 (June 2004).

60. Gao, S., Ver Steeg, G. & Galstyan, A. Efficient Estimation of Mutual Information for Strongly Dependent Variables. arXiv: 1411.2003 [cs.IT] (Nov. 2014).

61. Hodge, R. D., et al. Conserved cell types with divergent features in human versus mouse cortex. en. Nature 573, 61–68 (Sept. 2019).

62. Thul, P. J. & Lindskog, C. The human protein atlas: A spatial map of the human proteome. en. Protein Sci. 27, 233–244 (Jan. 2018).

63. MacParland, S. A., et al. Single cell RNA sequencing of human liver reveals distinct intrahepatic macrophage populations. en. Nat. Commun. 9, 4383 (Oct. 2018).

64. Uhlén, M. et al. Proteomics. Tissue-based map of the human proteome. en. Science 347, 1260419 (Jan. 2015).

65. GTEx Consortium. The GTEx Consortium atlas of genetic regulatory effects across human tissues. en. Science 369, 1318–1330 (Sept. 2020).

66. Gershman, A., et al. Epigenetic patterns in a complete human genome. en. Science 376, eabj5089 (Apr. 2022).

67. Kanehisa, M., Sato, Y., Kawashima, M., Furumichi, M. & Tanabe, M. KEGG as a reference resource for gene and protein annotation. en. Nucleic Acids Res. 44, D457–62 (Jan. 2016).

68. Gillespie, M., et al. The reactome pathway knowledgebase 2022. en. Nucleic Acids Res. 50, D687–D692 (Jan. 2022).

69. Slenter, D. N., et al. WikiPathways: a multifaceted pathway database bridging metabolomics to other omics research. en. Nucleic Acids Res. 46, D661–D667 (Jan. 2018).

70. Ocone, A., Haghverdi, L., Mueller, N. S. & Theis, F. J. Reconstructing gene regulatory dynamics from high-dimensional single-cell snapshot data. en. Bioinformatics 31, i89–96 (June 2015).

71. Qi, J., Sun, L., Li, K. & Wang, L. Gaussian noise parameter estimation based on multiple singular value decomposition and non-linear fitting. en. IET Image Proc. 16, 3025–3038 (Sept. 2022).

72. Bredikhin, D., Kats, I. & Stegle, O. MUON: multimodal omics analysis framework. en. Genome Biol. 23, 42 (Feb. 2022).

73. Selvaraj, S. & Natarajan, J. Microarray data analysis and mining tools. en. Bioinformation 6, 95–99 (Apr. 2011).

74. Johnson, W. E., Li, C. & Rabinovic, A. Adjusting batch effects in microarray expression data using empirical Bayes methods. en. Biostatistics 8, 118–127 (Jan. 2007).

75. Traag, V. A., Waltman, L. & van Eck, N. J. From Louvain to Leiden: guaranteeing well-connected commu- nities. en. Sci. Rep. 9, 5233 (Mar. 2019).

76. Wolf, F. A., et al. PAGA: graph abstraction reconciles clustering with trajectory inference through a topology preserving map of single cells. en. Genome Biol. 20, 59 (Mar. 2019).

77. McKenzie, A. T., et al. Brain Cell Type Specific Gene Expression and Co-expression Network Architectures. en. Sci. Rep. 8, 8868 (June 2018).

78. Whitfield, M. L., et al. Identification of genes periodically expressed in the human cell cycle and their expression in tumors. en. Mol. Biol. Cell 13, 1977–2000 (June 2002).

79. Stuart, T., et al. Comprehensive Integration of Single-Cell Data. en. Cell 177, 1888–1902.e21 (June 2019).

80. Chen, R. T. Q., Rubanova, Y., Bettencourt, J. & Duvenaud, D. Neural ordinary differential equations. arXiv: 1806.07366 [cs.LG] (June 2018).

81. Grathwohl, W., Chen, R. T. Q., Bettencourt, J., Sutskever, I. & Duvenaud, D. FFJORD: Free-form Con- tinuous Dynamics for Scalable Reversible Generative Models. arXiv: 1810.01367 [cs.LG] (Oct. 2018).

82. Shampine Lawrence, f. Some practical Runge-Kutta formulas. Math. Comput. 46, 135–150 (Jan. 1986).

83. Oliva, J. B. et al. Transformation Autoregressive Networks. arXiv: 1801.09819 [stat.ML] (Jan. 2018).

84. Friedman, J., Hastie, T. & Tibshirani, R. Sparse inverse covariance estimation with the graphical lasso. en. Biostatistics 9, 432–441 (July 2008).

85. Ledoit, O. & Wolf, M. A well-conditioned estimator for large-dimensional covariance matrices. J. Multivar. Anal. 88, 365–411 (Feb. 2004).

86. Beskos, A., Pillai, N. S., Roberts, G. O., Sanz-Serna, J. M. & Stuart, A. M. Optimal tuning of the Hybrid Monte-Carlo Algorithm. arXiv: 1001.4460 [math.PR] (Jan. 2010).

87. Neal, R. M. MCMC using Hamiltonian dynamics. arXiv: 1206.1901 [stat.CO] (June 2012).

88. *tfp.mcmc.HamiltonianMonteCarlo* en. https://www.tensorflow.org/probability/api_docs/python/tfp/mcmc/HamiltonianMonteCarlo. Accessed: 2023-4-18.

89. Pedregosa, F., et al. Scikit-learn: Machine Learning in Python. J. Mach. Learn. Res. 12, 2825–2830 (2011).

90. Lobo, J. M., Jiménez-Valverde, A. & Real, R. AUC: a misleading measure of the performance of predictive distribution models. en. Glob. Ecol. Biogeogr. 17, 145–151 (Mar. 2008).

91. Flach, P. A., Hernández-Orallo, J. & Ramirez, C. F. *A Coherent Interpretation of AUC as a Measure of Aggregated Classification Performance* Jan. 2011.

92. Fawcett, T. An introduction to ROC analysis. Pattern Recognit. Lett. 27, 861–874 (June 2006).

93. Calders, T. & Jaroszewicz, S. *Efficient AUC Optimization for Classification* in *Knowledge Discovery in Databases: PKDD* 2007 (Springer Berlin Heidelberg, 2007), 42–53.

94. Jurman, G., Visintainer, R., Filosi, M., Riccadonna, S. & Furlanello, C. The HIM glocal metric and kernel for network comparison and classification in 2015 IEEE International Conference on Data Science and Advanced Analytics (DSAA) (Oct. 2015), 1–10.

